# Unbiased Single-Cell Transcriptome-Proteome Co-Profiling Reveals Malignant Dormancy and Post-Transcriptional Buffering of CTCs

**DOI:** 10.64898/2026.03.17.711781

**Authors:** Liyong He, Kaiqiang Ye, Huiying Li, Lin Jiang, Wenyi Zhang, Kaitong Dang, Xiaoying Ma, Jiahao Shen, Yuwei Dong, Wenjia Wang, Handong Wang, Zhen-Li Huang, Yan Huang, Nan Xiang, Zhenyu Yin, Xiangwei Zhao

## Abstract

Leptomeningeal metastasis is driven by rare cerebrospinal fluid circulating tumor cells (CSF-CTCs). However, the mechanisms underlying their adaptation to chemotherapeutic stress remain elusive, primarily because transcriptomics alone poorly predicts functional protein states. Here, we present scMAPS, a single-cell multi-omics method that employs magnetic-assisted partitioning cell lysates to enable unbiased transcriptome-proteome co-profiling without loss-prone physical splitting and precision device. By coupling scMAPS with our custom CLEAP (CTC Label-free Enrichment and Accurate Picking) system, we performed the first deep multi-omics profiling of rare clinical CSF-CTCs before and after localized chemotherapy, detecting an average of 2,547 proteins and 7,821 genes per cell. The integrated CLEAP-scMAPS pipeline reveals a coordinated prioritizing survival over proliferation malignant dormancy phenotype and identified post-transcriptional buffering as the primary driver of treatment resistance. Our platform enables the comprehensive molecular phenotyping of rare clinical specimens, providing a highly versatile framework for decoding complex post-transcriptional regulatory networks.

## Introduction

Leptomeningeal metastasis (LM), a lethal complication of advanced lung cancer, is primarily driven by cerebrospinal fluid circulating tumor cells (CSF-CTCs)^1–3^. Mechanistically dissecting how these extremely rare cells evade therapeutic stress remains a formidable clinical challenge. Addressing this requires overcoming two fundamental hurdles: the intact isolation of individual CTCs from complex biofluids (cutoff for diagnosis: ≥1 cell/mL^4^) and the unbiased simultaneous profiling of their multi-omics landscapes. Conventional bulk analyses mask critical cellular heterogeneity, whereas droplet-based and FACS platforms are incompatible with such extreme sample sparsity. Furthermore, transcriptome profiling alone (scRNA-seq)^5, 6^ alone poorly predicts the abundance of functional proteins, the ultimate effectors of cellular phenotypes^7^, due to pervasive post-transcriptional regulation and complex proteostasis^8, 9^. Consequently, simultaneous transcriptome and proteome profiling of the same individual cell is imperative^10, 11^.

Existing techniques remain inadequate for this dual requirement. Conventional cell-sorting technologies fail to recover rare cells at high-purity without relying on predefined surface markers, thereby introducing severe selection bias. Regarding single-cell multi-omics (scMulti-omics) profiling, current antibody-dependent methods, such as CITE-seq^12^ and Prox-seq^13^, are constrained by target availability, precluding unbiased and proteome-wide discovery^14^. Conversely, integrating unbiased mass spectrometry (MS)^15–17^ with scRNA-seq offers a promising route but typically requires the physical splitting of nanoliter lysates^18, 19^, which inherently risks the dilution and loss of ultra-low-abundance molecules. Moreover, these platforms demand expensive nanoliter dispensing robots or specialized microfluidic devices that impede widespread adoption in standard laboratories.

To overcome these limitations, we engineered a fully integrated pipeline that couples our custom-built CTC Label-free Enrichment and Accurate Picking (CLEAP) system with a novel single-cell Magnetic-Assisted Partitioning Sequencing (scMAPS) strategy. CLEAP resolves the isolation challenge by ensuring the intact, marker-free recovery of individual CTCs. To circumvent the sample loss inherent to conventional physical splitting, scMAPS employs streptavidin microbeads to chemically partition lysates, efficiently capturing RNA for multiplexed full-length transcriptome sequencing^20^ while preserving the intact protein supernatant for in-tip shotgun-MS^21^. Using this robust and device-free platform, we performed the first unbiased transcriptome-proteome co-profiling of rare clinical CSF-CTCs. Notably, the integrated CLEAP-scMAPS pipeline revealed an adaptive shift prioritizing survival over proliferation, alongside extensive post-transcriptional buffering, to navigate localized therapeutic stress. Ultimately, this approach provides a highly versatile framework for the comprehensive molecular phenotyping of scarce clinical specimens.

## Results

### Establishment and characterization of the CLEAP system for pristine isolation of rare single CSF-CTCs

To decode the molecular vulnerabilities of LM at single-cell resolution, we engineered an integrated upfront physical isolation pipeline, termed the CTC Label-free Enrichment and Accurate Picking (CLEAP) system, for the pristine isolation of rare CSF-CTCs (Fig. 1). Within this system, clinical CSF collected via lunger cancer patient Ommaya reservoirs^1^ is first subjected to a slanted spiral microfluidic chip^22^ for rapid and label-free CTCs enrichment. This step is seamlessly coupled with a custom-built capillary microneedle sampling platform^23, 24^ to visually confirm and precisely isolated individual CTCs into microtubes. Integrating the stringent physical capture of the CLEAP system with our downstream scMAPS method enables the simultaneous, unbiased measurement of the transcriptome and proteome within the same single-cell, providing a robust analytical framework for downstream mechanistic discovery.

**Fig. 1.**
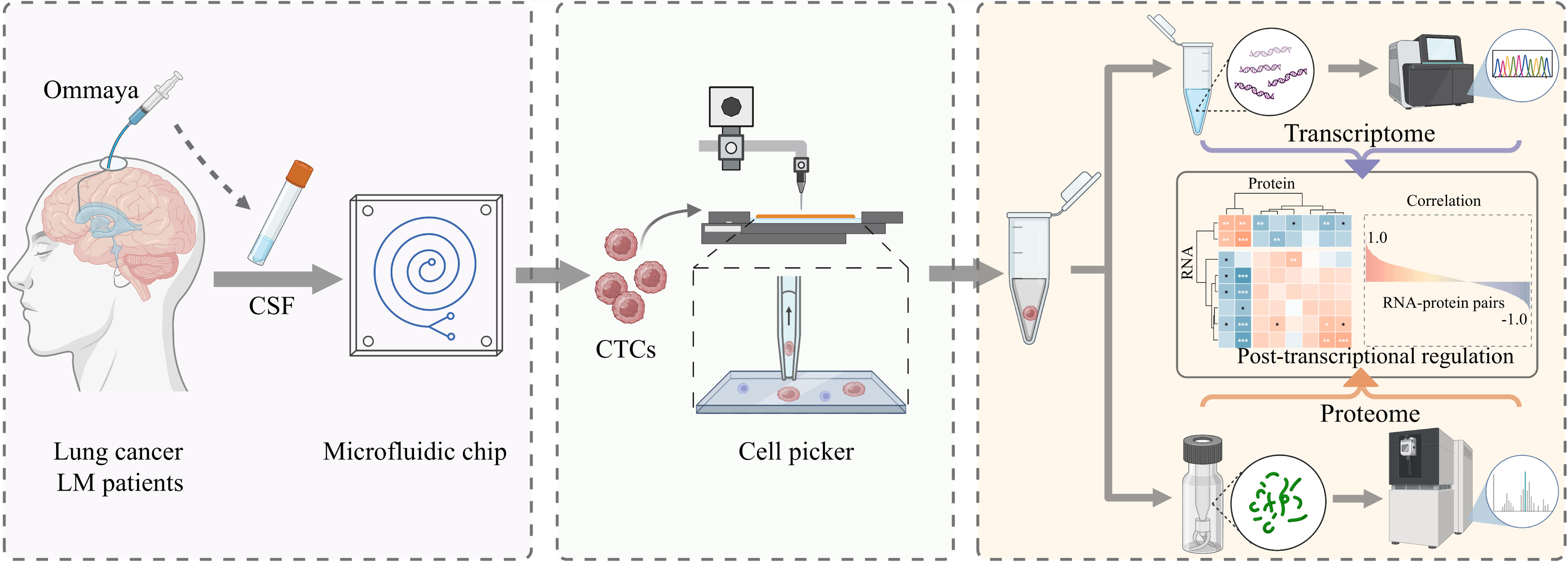
Schematic overview of the integrated CLEAP workflow for single-cell multi-omics analysis of rare CSF-CTCs.

To rigorously evaluate the two-stage physical isolation capabilities of the CLEAP system, we first characterized the upfront module, which features a polymer microfluidic chip with slanted spiral channels (Fig. 2a, b). This device achieves rapid, label-free target enrichment by exploiting size-dependent inertial focusing within a trapezoidal cross-section. Microscopic tracking at the channel outlet demonstrated a clear spatial divergence of these size-fractionated cell trajectories, confirming that the device efficiently isolates a highly enriched CTCs population with robust recovery rates from complex fluid mixtures (Fig. 2c).

**Fig. 2.**
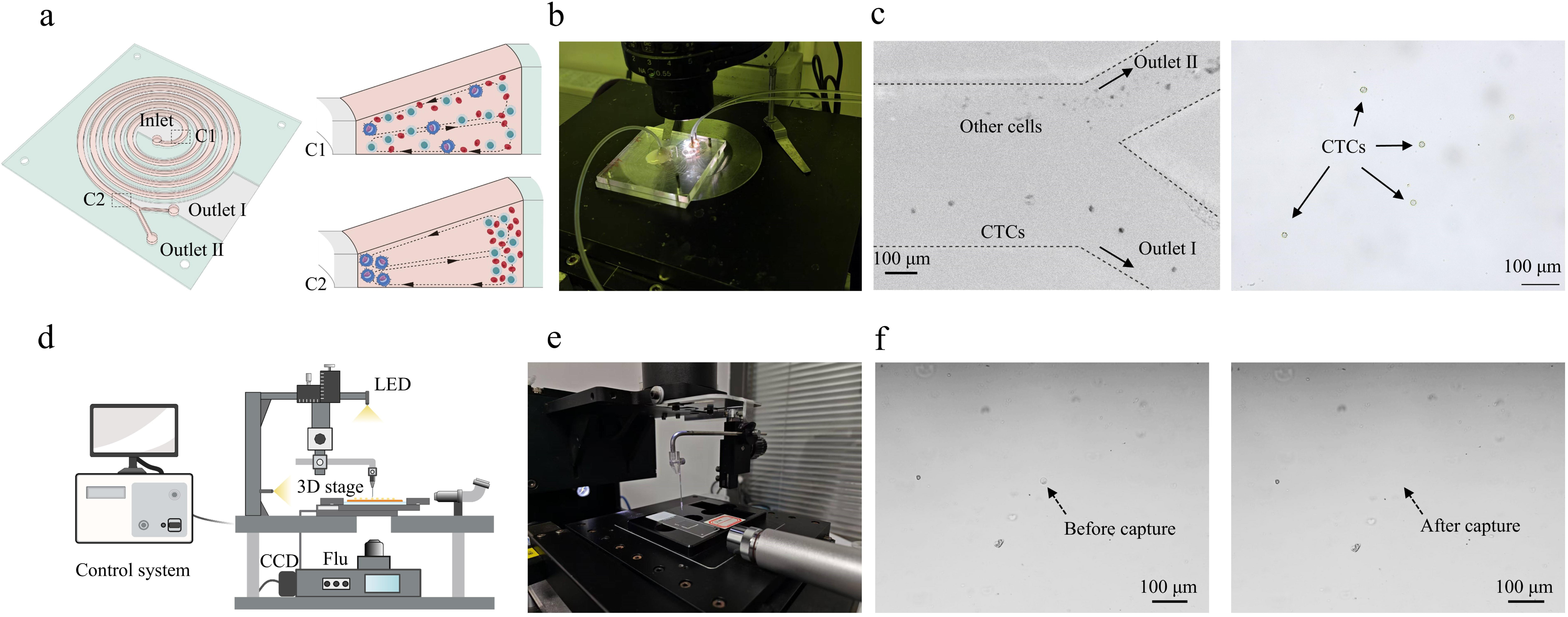
CLEAP system for pristine isolation of rare single CSF-CTCs. Schematic diagram of the polymer microfluidic chip with slanted spiral channels (a) and photograph of the fabricated device (b). c, Microscopy images illustrating the distributions of CTCs cells and other cells at the microfluidic channel outlet (left) and enriched CTCs population (right). Side view schematic diagram of the custom-built capillary microneedle sampling platform (d) and photograph of the instrument (e). f, Microscopic bright images of individual CTCs before (left) and after (right) capture on the microneedle sampling platform.

To bridge the gap between this bulk-enriched suspension and the stringent purity requirements of single-cell profiling, we subsequently employed a precision-engineered capillary microneedle sampling platform (Fig. 2d, e). High-resolution bright-field microscopy confirmed the operational precision of this module, precisely capturing the exact spatial coordinates of individual CTCs immediately before and seamlessly after their targeted, non-destructive aspiration into the capillary tip (Fig. 2f). Applying this high-precision workflow to standard 10 mL clinical CSF samples achieved a robust yield ranging from ∼30 to upwards of 100 intact single CTCs. Altogether, these empirical characterizations demonstrate that the CLEAP system effectively overcomes the immense cellular background of clinical CSF, delivering pristine, intact single CTCs for downstream mechanistic discovery.

### Design of scMAPS method for simultaneous unbiased single-cell transcriptome-proteome co-profiling

To overcome the potential dilution and loss of ultra-low abundance molecules caused by physical sample-splitting strategies, we have designed a device-free, magnetic-assisted chemical partitioning strategy for simultaneous multi-omics co-profiling. The scMAPS method, illustrated in Fig. 3a, integrates single-cell lysis, streptavidin (SA) microbead-mediated molecular partitioning, and simultaneous transcriptome-proteome profiling. Briefly, the isolated individual mammalian cells were transferred into PCR tubes containing 2 μl of lysis buffer (0.1% n-Dodecyl-β-D-Maltoside (DDM) in 10 mM Tris (pH 8), supplemented with 0.5 μM of biotin-dT_25_ probe). Following a single freeze-thaw cycle and DDM-assisted incubation to release cellular RNA and proteins, the biotin-dT_25_ probes in the lysis buffer hybridize with poly(A^+^) RNA. Subsequently, SA-microbeads are introduced to the lysate to capture and enrich the poly(A^+^) RNA/biotin-dT_25_ heteroduplex. The lysate supernatant containing the proteins is transferred to C18-spintip device through magnetic separation for in-tip shotgun proteomics workflow and LC-MS/MS analysis, while the mRNA is eluted from SA-bead.s-coupled heteroduplex for multiplexed sequencing of full-length transcriptome analysis using our previously established CBTi-seq protocol^20^.

**Fig. 3.**
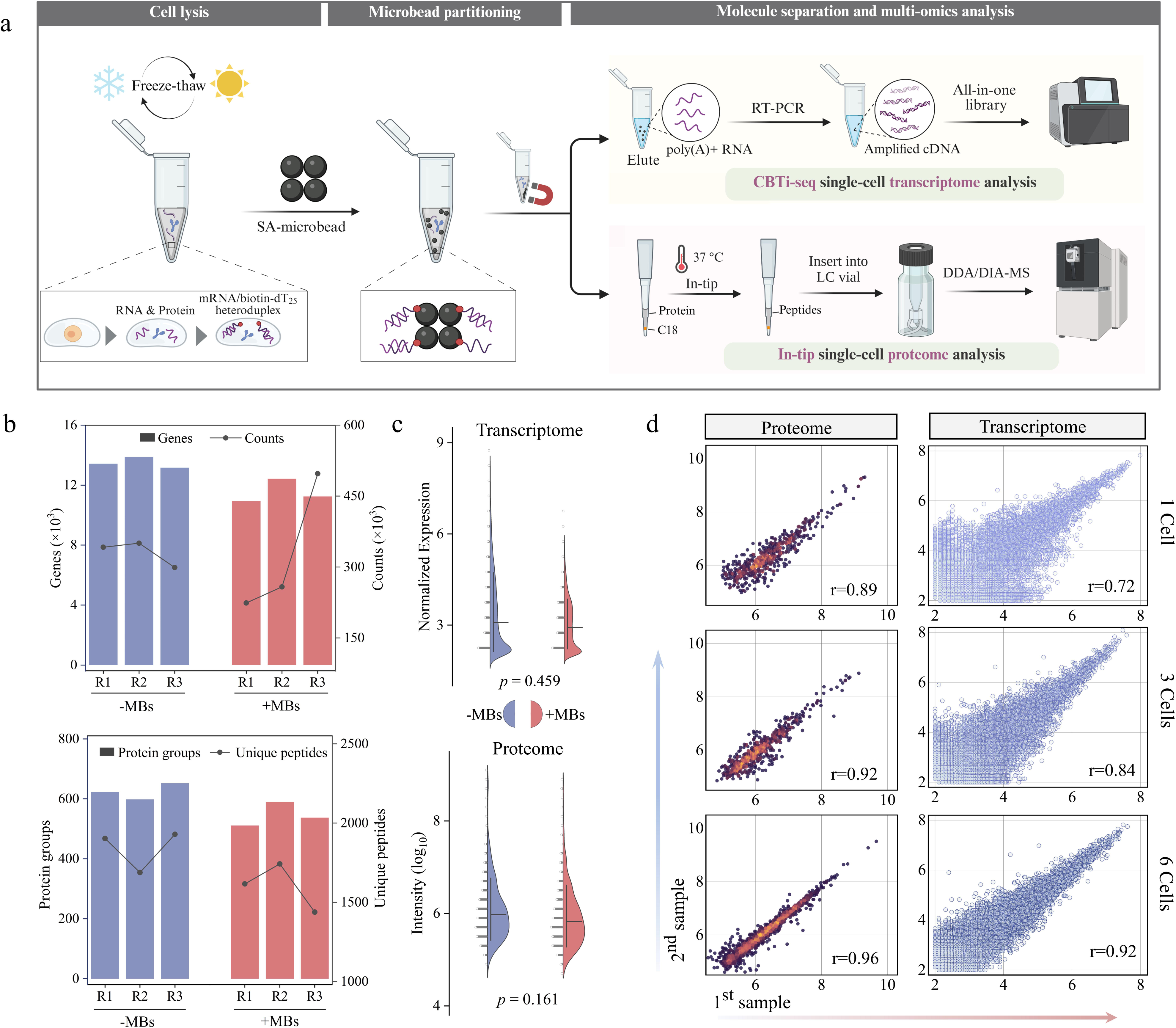
Schematic and performance of the scMAPS method for single cell multi-omics analysis. a, Overview of the workflow including cell lysis, microbead capture, and molecule separation for downstream CBTi-seq transcriptome and in-tip proteome co-profiling. b, Number of genes and counts identified for 30 pg of total RNA extracted from hESC cell lysates (n=3) without and with the addition of MBs (top panel); Number of protein groups and unique peptides identified for 0.6 ng of proteins from hESC cell lysates (n=3) without and with the addition of MBs (bottom panel). c, The normalized quantitative expression differences of genes (top panel) and proteins (bottom panel) identified without and with MBs. *p* values calculated using paired t test. d, Pairwise correlation of transformed gene and protein expressions among 1, 3 and 6-hESC cell samples (two biological replicates per condition).

Microscopic characterization confirmed the highly efficient capture of poly(A^+^) RNA/biotin-dT_25_ heteroduplexes by SA-beads (Supplementary Fig. 1). Crucially, unlike commercial oligo(dT) alternatives that suffer from severe non-specific protein adsorption, uniquely achieved an optimal balance of maximizing mRNA recovery while minimizing protein loss (Supplementary Fig. 2). We evaluated with ultra-low inputs (equivalent to 3 cells: a mixture of 600 pg protein and 30 pg RNA), the bead-based partitioning system (+MBs) achieved molecular identification depths (Fig. 3b) and quantitative profiles comparable to the bead-free control (-MBs) (paired t test: *p* > 0.05, n = 3, Fig. 3c). To evaluate scMAPS reproducibility, we isolated precise inputs of 1- to 6-hESC cell utilizing our custom-built capillary microneedle platform, with the capturing process visually verified under bright-field and fluorescence microscopy (Supplementary Fig. 3). We demonstrated robust quantitative reproducibility, with proteome correlations (r = 0.89–0.96) consistently outperforming transcriptome correlations (r = 0.72–0.92) (Fig. 3d).

To maximize proteomic sensitivity for trace-level inputs, we engineered a minimalistic sample preparation workflow that integrates DDM surface passivation, bypasses loss-prone chemical derivatization and desalting steps. This streamlined processing was subsequently executed in tandem with tailored data dependent acquisition (DDA)-MS acquisition parameters (ITmax: 300 ms, IW: 4 Th) (Supplementary Fig. 4). Crucially, benchmarking against physical sample-splitting technique (where half the lysate is used for transcriptome and half for proteome) demonstrated that the scMAPS chemical partitioning strategy effectively circumvents dilution-induced losses, achieving molecular identification depths comparable to intact, unpartitioned single cells (Supplementary Fig. 5).

### Accuracy, scalability and superiority for scMAPS

Prior to profiling scarce clinical specimens, we systematically benchmarked the analytical sensitivity, quantitative accuracy, and scalability of scMAPS using standard cell lines. We first validated the independent performance of multiplexing transcriptome full-length sequencing, CBTi-seq, across varying multiplexing levels (from 4- to 32-plex) in hESC cells. The analysis demonstrated that gene detection sensitivity and inter-sample quantitative correlations remained highly stable as cellular throughput increased (Fig. 4a, Supplementary Fig. 6). Furthermore, multiplexed CBTi-seq maintained uniform full-length gene body coverage without significant 3’ or 5’ bias (Fig. 4b). To rigorously benchmark this approach, we compared CBTi-seq against the gold-standard SmartSeq2 protocol^25^ using L929 and hESC cell lines. CBTi-seq achieved exceptional detection depth and exhibited high concordance with SmartSeq2 in overall gene identification (Fig. 4c, Supplementary Fig. 7). Notably, while both methods yielded comparable read mapping rates, CBTi-seq displayed superior coverage uniformity across the gene body (Fig. 4d). This complete read distribution pattern was visually corroborated by Integrated Genome Viewer (IGV) tracking of representative transcripts, such as POU5F1 and Col1a1 (Fig. 4e).

**Fig. 4.**
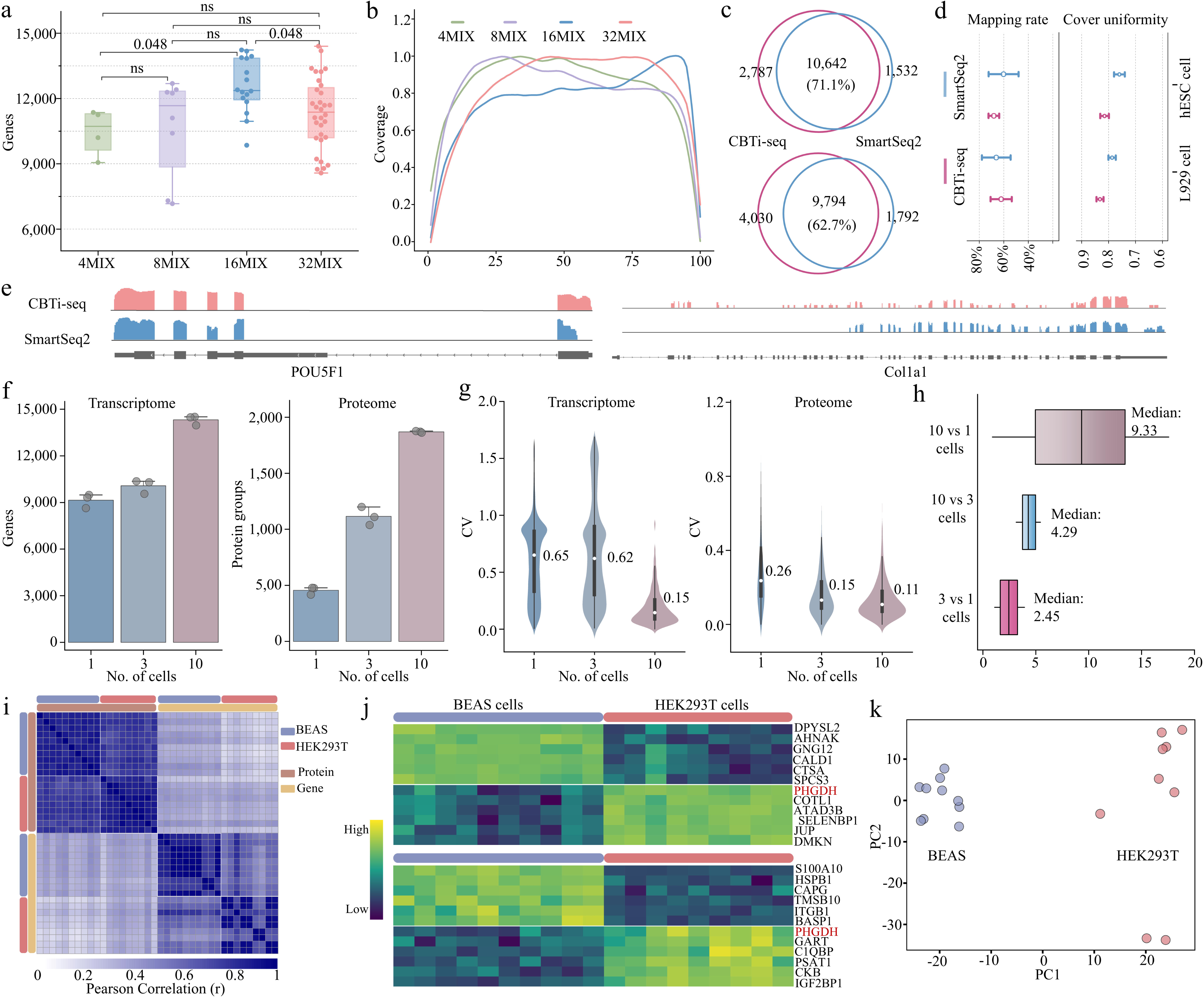
Analytical benchmarking, scalability, and cell-type classification of the scMAPS method. The average gene detection (a) and gene body coverage (b) performance obtained through sequencing under the conditions of different multiplexed hESC cells (4, 8, 16, and 32MIX). Statistical differences were analyzed using a two-sided Wilcoxon rank-sum test. Adjusted p-values are presented above boxplots; ns, not significant. c, Venn diagram of the identified genes by CBTi-seq and Smart-seq2 in the three replicates of L929 cells (top panel) and hESC cells (bottom panel). d, The percentage of mapping reads and coverage uniformity across gene body for each method. e, IGV tracks showed the coverage of two representative transcripts (POU5F1 in hESC cells and Col1a1 in L929 cells) using CBTi-seq and Smart-seq2. f, The average number of detected genes (left) and protein groups (right) in 1, 3, and 10 HEK293T cells (protein groups were detected under the DDA mode). Error bars indicate standard deviations (± s.d.). g, Distributions of the coefficients of variation (CV) for quantified genes (left) and proteins (right) between the different pooled cells. Indicated values represent median CVs (center point in the diagram). h, The quantitative protein abundance ratio between the median intensities in 10, 3, and 1 cell. The central lines in the boxes indicate the median values. Boxplot boundaries extend from the first (25%) and third (75%) quartile, representing the interquartile range. i, Pearson correlation heatmap with clustering of transcriptome and proteome results for both single BEAS (n = 10) and HEK293T (n = 9) cells (protein groups are detected under the DIA mode). j, Top 6 differentially expressed genes from transcriptome (top panel) and proteins from proteome (bottom panel) for each cell type. Candidate marker features were determined using a two-sided Wilcoxon rank-sum test (*p.adj* < 0.001). k, PCA cluster of BEAS and HEK293T cells for the multi-omics joint analysis.

Building upon the robust performance of transcriptomic module, we evaluated the multi-omics scalability of the integrated scMAPS workflow across ultra-low inputs of 1, 3, and 10 HEK293T cells. The platform exhibited highly proportional molecular scaling; notably, identification depths expanded from an average of 9,148 genes and 457 protein groups at the single-cell to over 14,300 genes and 1,870 proteins in 10-cell populations (Fig. 4f). Further, Venn diagrams confirmed excellent technical reproducibility and consistent compositional overlap across both modalities (Supplementary Fig. 8). MS quality control metrics for the 3-cell samples further attested to the reliability and stability of the proteome data (Supplementary Fig. 9). For quantitative accuracy, as expected, median coefficients of variation (CVs) progressively decreased with increasing cell inputs due to the averaging of underlying cell-to-cell stochasticity (Fig. 4g). Notably, we observed significantly higher CVs for the transcriptome compared to proteome, a finding aligned with previous reports^16, 26^. This biological phenomenon elegantly reflects the buffering of short-lived transcriptional “bursts”, whereas proteins generally possess longer half-lives and higher copy numbers, buffering expression noise^27^. Quantitative accuracy was visually supported by robust normalized TPM distributions (Supplementary Fig. 10) and orthogonally validated using the “Proteomic Ruler” method^28^, which accurately recapitulated the theoretical abundance ratios of 35 histone-related proteins across varying cell inputs (Fig. 4h). Crucially, scMAPS captured an expansive proteomic dynamic range spanning 4∼5 orders of magnitude, achieving unbiased subcellular coverage that enables the robust detection of intracellular and nuclear regulatory proteins typically inaccessible to surface-marker-dependent technologies (Supplementary Fig. 11).

To demonstrate the ultimate biological utility of scMAPS in cell-type classification, we coupled the workflow with a data-independent acquisition (DIA)-based Orbitrap Astral MS, which significantly outperformed conventional DDA model in trace-level sensitivity (Supplementary Fig. 12). Profiling BEAS-2B and HEK293T cells yielded deep multi-omic signatures (averaging >9,000 genes and >1,200 proteins per cell, Supplementary Fig. 13a) that distinctly separated the two lineages based on Pearson correlation clustering (Fig. 4i). Qualitatively, nearly all detected proteins possessed corresponding mRNA transcripts (Supplementary Fig. 14c, d), with proteome signals predominantly originating from high-abundance mRNAs (Supplementary Fig. 13b). However, despite this extensive compositional overlap, intracell cross-modality quantitative correlations remained modest (r = 0.2-0.45) (Supplementary Fig. 14b). The sharp contrast highlights the stringent and non-linear translation dynamics, and this inherent multi-omics discordance critically impacts cell-type characterization. When extracting the top 6 significantly enriched markers for each lineage, we observed minimal overlap between transcriptome and proteome data (Fig. 4j), demonstrating that widely used scRNA-seq is insufficient to accurately predict functional protein markers. Consequently, while single-modality principal component analysis (PCA) yielded suboptimal separation for certain cells (Supplementary Fig. 15), joint dimensional reduction of the combined multi-omics matrix completely resolved these ambiguities. This integrated approach achieved highly precise cell-type clustering (Fig. 4k) and enabled the robust cross-validation of shared dual-omics markers, such as PHGDH (Supplementary Fig. 16), establishing scMAPS as a superior tool for accurate cell type classification and marker validation.

### Multi-omics profiling reveals coordinated immune evasion and metabolic reprogramming in chemotherapy-surviving CSF-CTCs

Equipped with this robust and highly sensitive multi-omics engine, we returned to the clinically challenging CSF-CTCs to dissect their adaptive responses to localized chemotherapy within the CSF microenvironment. Specifically, we obtained CSF samples from lung cancer patients before and after intracranial pemetrexed administration, isolating individual CSF-CTCs via our CLEAP system. Robust transcriptome expression of canonical tumor markers unambiguously verified the malignant identity and high purity of the captured single cells (Fig. 5a). Using the scMAPS workflow, an average of 2,547 proteins and 7,821 genes identified in control CSF-CTCs (15-25 μm, n=13) as well as 2,514 proteins and 6,170 genes for treated CSF-CTCs (15-25 μm, n=12) (Supplementary Fig. 17). Integrated clustering revealed a profound state shift between treatment-naive and surviving CTC populations (Fig. 5b). Gene Set Enrichment Analysis (GSEA) unveiled a striking and coordinated downregulation of immune pathways at both transcriptome and proteome levels including interferon-γ (IFN-γ) response, complement system, and inflammatory (Fig. 5c, d; Supplementary Fig. 18), establishing an immune-cold phenotype. Specifically, core complement components (C3^29^, C1S/R) were suppressed (Fig. 5e; Supplementary Fig. 19), likely dampening complement-dependent cytotoxicity. Notably, STAT1 and LAP3 exhibited RNA-protein discordance within the IFN-γ pathway^30, 31^ (Fig. 5f), suggesting post-transcriptional buffering in fine-tuning this adaptive response.

**Fig. 5.**
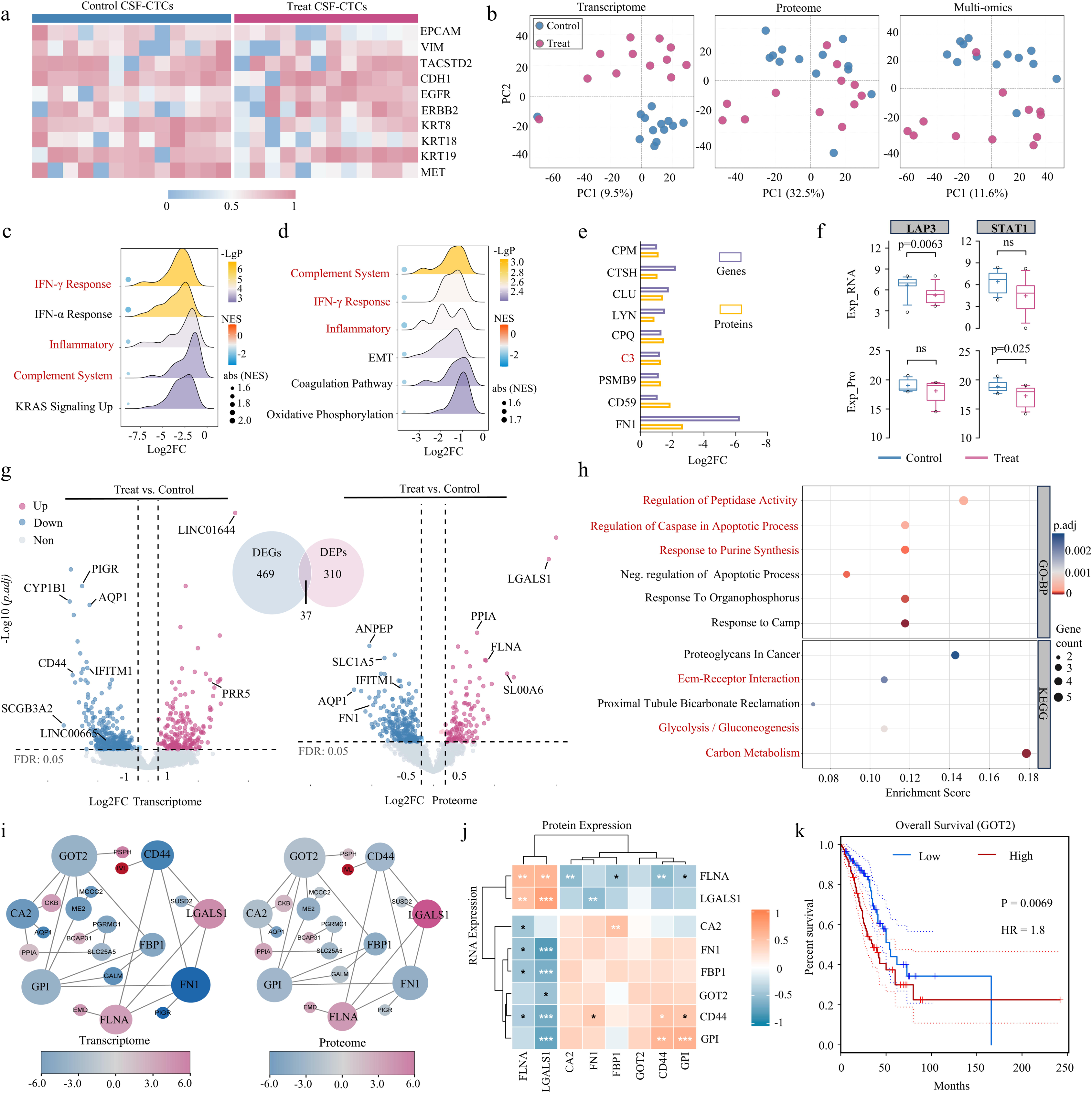
Multi-omics profiling of CSF-CTCs in pemetrexed-treated leptomeningeal metastasis. a, The gene expression heatmap of common tumor markers in CSF-CTCs. b, PCA of control CTCs and treat CTCs groups based on the expression matrix of transcriptome (left), proteome (middle) and multi-omics joint analysis (right). GSEA ridge plot displaying transcriptome (c) and proteome (d) pathways differentially enriched in treat vs. control groups. e, Co-downregulated genes in the complement pathway based on log_2_FC which identified across both transcriptome and proteome. f, Quantitative comparison of LAP3, and STAT1 expression at mRNA and protein levels. *p*-values are obtained from paired two-tailed test. ns, no significant. The boxes indicate the median value, interquartile range, with whiskers extending from the box boundaries to upper/lower quartile ± 1.5 interquartile range. g, Volcano plots of differential gene (left) and protein (right) expression, with an overlapping Venn diagram of differentially expressed genes (DEGs) and proteins (DEPs). DEGs: (|FC| > 2, p.adj < 0.05); DEPs: (|FC| > 1.5, p.adj < 0.05). h, GO-Biological Process (GO-BP) and KEGG pathway enrichment analysis of overlapping DEGs/DEPs. i, Protein-protein interaction (PPI) networks of overlapping DEGs/DEPs, with red indicating upregulation and blue indicating downregulation in treat vs. control groups. j, Correlation analysis of key genes across transcriptome and proteome features, highlighting consistency in expression changes. k, Kaplan-Meier overall survival analysis of the metabolic regulator GOT2 in the TCGA-LUAD cohort (n = 120). TCGA, The Cancer Genome Atlas.

To pinpoint the core molecular drivers of this resistance, we performed differential expression analysis, identifying 506 genes and 347 proteins that were significantly altered post-treatment (Fig. 5g, Supplementary Fig. 20). Notably, canonical tumor markers (SCGB3A2^32^, SFTPB^33^, and CD44^34^) were all significantly downregulated post-treatment. Intersecting these datasets revealed 37 core targets that were consistently dysregulated across both modalities (Fig. 5g, Supplementary Table 1). Functional enrichment (Fig. 5h) molecularly validated pemetrexed efficacy via suppressed purine synthesis. Crucially, the concurrent downregulation of caspase regulators implies a release of the apoptotic brake, a state further corroborated by the marked 3’ transcripts bias (Supplementary Fig. 21) indicative of caspase-mediated RNA degradation. In parallel, KEGG analysis highlighted suppressed cell adhesion and metabolic rewiring, consistent with a transition towards dormancy. Within PPI networks (Fig. 5i), ECM-remodeling hubs (CD44, FN1^35^) (Supplementary Fig. 22) and metabolic regulators (FBP1, GPI, GOT2) (Supplementary Fig. 23 and 24) formed distinct functional clusters. Crucially, cross-omics analysis revealed a positive correlation between these adhesive and metabolic machineries (Fig. 5j), confirming their synchronized suppression as a deliberate survival program.

Clinical validation in the TCGA-LUAD cohort confirmed that high expression of the anabolic drivers GOT2^36, 37^/GPI^38^ (Fig. 5k, Supplementary Fig. 25a) and low expression of the tumor suppressor FBP1^39^ are individually associated with poor survival (Supplementary Fig. 25b). Strikingly, within the therapeutic CSF microenvironment, we observed a synchronized downregulation of both GOT2/GPI and FBP1. In contrast to the Warburg effect where FBP1 loss disinhibits glycolysis to fuel proliferation, its depletion in these treated CSF-CTCs coincided with a broad suppression of glycolytic pathway. This concurrent suppression defines a distinct state of malignant dormancy, revealing a survival-over-growth adaptive strategy wherein CSF-CTCs compromise global metabolic activity to withstand chemotherapeutic pressure.

### Integrated proteogenomic multi-omics analysis reveals post-transcriptional regulatory modes driving therapeutic adaptation

We next leveraged the matched multi-omics data to dissect the regulatory logic of CSF-CTCs. The scMAPS workflow achieved high proteogenomic coverage, with 88.7% of the total detected proteins (n = 5,079) having corresponding transcripts identified (Fig. 6a). Consistent with the non-linear “central dogma” observed in bulk proteogenomics, global correlation analysis of transcript-protein pairs with <50% missing values revealed a modest overall correlation (R = 0.2335), highlighting the pervasive influence of post-transcriptional buffering in maintaining cellular homeostasis (Fig. 6b). Further individual correlation analysis of 1,307 quantified gene-protein pairs (Fig. 6c) identified 28 positively correlated pairs (r > 0.5), primarily involved in the regulation of translation, signal transduction, and RNA metabolic processes (Supplementary Fig. 26a). Notably, 20 negatively correlated pairs (r < −0.4) were enriched in RNA splicing pathways (Supplementary Fig. 26b). Among them, SNRPA1^40^ as a pivotal splicing regulator, was characterized by a treatment-specific discordance between RNA and protein levels. Sashimi plot visualization (Fig. 6d) mechanistically linked this protein depletion to a drug-induced shift toward non-productive splicing isoforms targeted by nonsense-mediated decay (AS-NMD). Crucially, this perturbation propagated to downstream oncogenic splicing networks. We observed a concurrent suppression of *PLEC* exon 31 inclusion (Fig. Supplementary Fig. 27), an SNRPA1-dependent event essential for invasion and metastasis^40^. Collectively, these data reposition SNRPA1 from a generic spliceosomal component to a critical effector whose post-transcriptional silencing drives treatment-mediated anti-metastatic effect. This autoregulatory layer highlights specific splicing factors (SNRPA1, SRSF6^41^, PRPF8^42^) as stealth therapeutic targets invisible to transcriptomics alone^43, 44^.

**Fig. 6.**
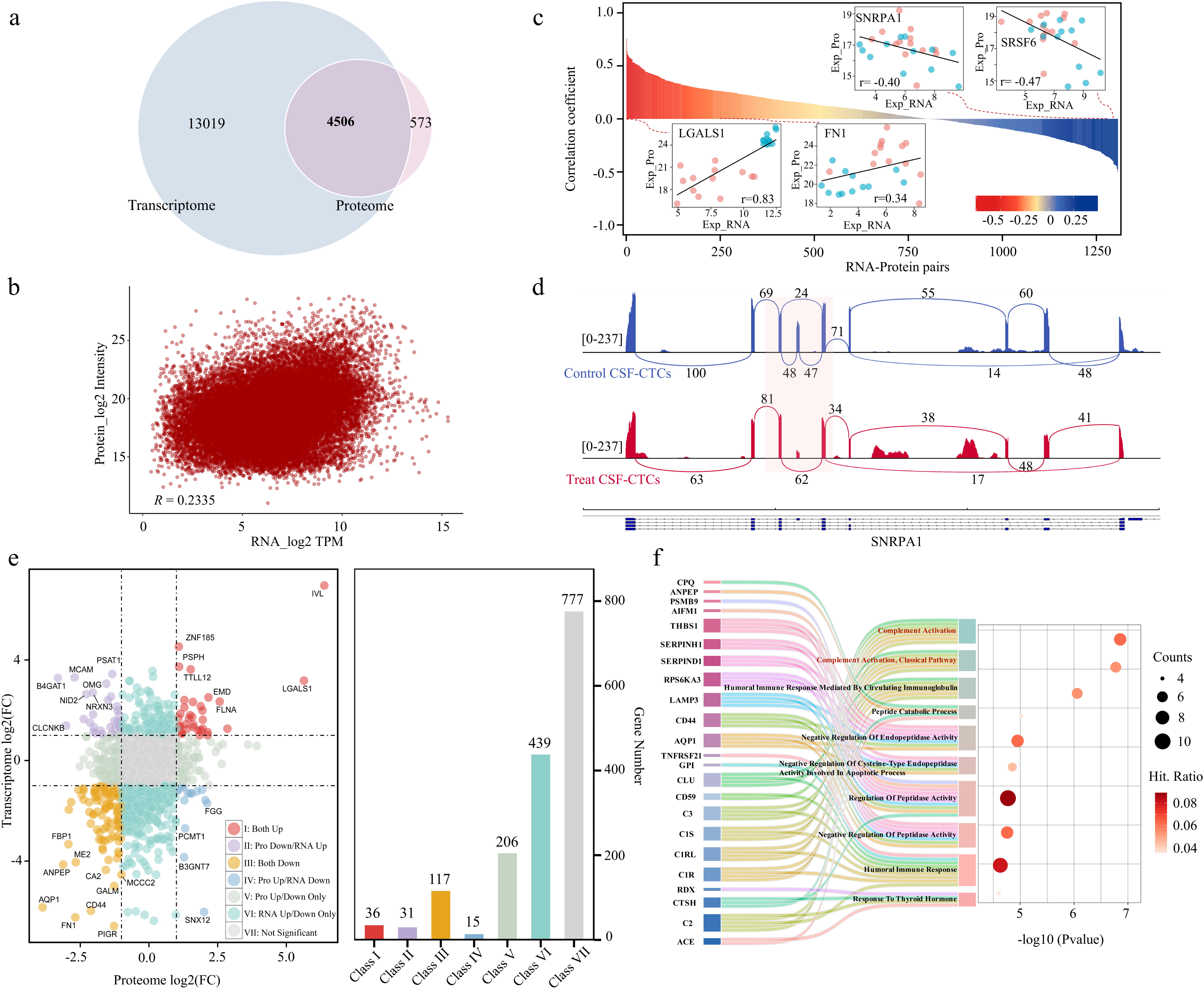
Correlation analysis of transcriptome and proteome in CSF-CTCs. a, Venn diagram of total identified genes in transcriptome and proteins in proteome in the CSF-CTCs. b, Correlation analysis of total transcript-protein pairs in CSF-CTCs (R = 0.2335, n = 1,307). c, Correlation distribution of the transcript-protein pairs in CSF-CTCs (n = 1,307). d, Treatment-induced aberrant splicing of SNRPA1. Sashimi plots visualization of RNA-seq reads mapping to the SNRPA1 locus in control (top, red) versus treat (bottom, blue) CTCs. Loops represent splice junctions, and the height of bars represents read coverage. e, Two-dimensional scatter plot of the transcript-protein pairs in (C) (n = 1,307). Class I: up-regulated in both RNA and protein; Class II: up-regulated in RNA while down-regulated in protein; class III: down-regulated in both RNA and protein; class IV: up-regulated in protein while down-regulated in RNA; class V: protein up/down-regulated only. class VI: RNA up/down-regulated only. class VII: no significantly different in both RNA and protein. f, Gene ontology term enrichment analysis (p.adj < 0.05) of class III genes (orange) in (e), including the down-regulated genes in both transcriptome and proteome.

Furthermore, association analysis demonstrated significant discordance between mRNA and protein-level differential changes, leading us to categorize genes into seven distinct regulatory classes (Fig. 6e). Among these, Class III genes (concomitant downregulation) were significantly enriched in the complement pathway (Fig. 6f), and were consistently verified in KEGG analysis (Supplementary Fig. 28), revealing an immune evasion mechanism within the CSF microenvironment. In contrast, Class II genes (RNA-up, protein-down) were enriched in wound healing and coagulation pathways (Supplementary Fig. 29a). This discordance highlights a frustrated repair phenotype, while chemotherapy-induced damage triggers a transcriptional compensation response, translational bottlenecks or rapid turnover prevent the accumulation of functional repair proteins. Moreover, we observed that the low global correlation was largely driven by Class VI (using data on RNA-down, protein-stable). Functional enrichment of this class in purine metabolism and the TCA cycle points to a metabolic buffering strategy (Supplementary Fig. 29b), where stable enzyme pools persist to sustain basal bioenergetics despite the widespread transcriptional shutdown induced by pemetrexed. This prevalence of post-transcriptional buffering^45^ further underscores the necessity of direct proteome profiling to accurately capture the functional state of drug-tolerant cells.

## Discussion

The grand challenge in deciphering therapeutic resistance in rare clinical specimens, such as CSF-CTCs in leptomeningeal metastasis, lies in the dual bottleneck of achieving pristine cellular isolation and depth of multi-omics profiling. Current methodologies often falter on both fronts, relying on biased, affinity-based enrichment that compromises cellular recovery, and downstream physical lysate splitting that dilutes ultra-low-abundance molecules. In this article, we establish a fully integrated end-to-end pipeline that seamlessly couples our custom CLEAP system with the scMAPS strategy. By synergizing label-free microfluidic enrichment and microneedle-based deterministic isolation with magnetic-assisted chemical partitioning, this robust workflow eliminates both the background noise of complex biofluids and the manipulative sample loss associated with physical splitting. Ultimately, this integrated approach democratizes single-cell proteogenomics, empowering standard biomedical laboratories to perform deep, unbiased transcriptome-proteome co-profiling of scarce liquid biopsies without the reliance on complex nanoliter-dispensing automation.

Deploying the CLEAP-scMAPS pipeline on the clinical frontlines immediately unlocked biological dimensions invisible to single-modality approaches. By unbiasedly co-profiling thousands of genes and proteins in individual CSF-CTCs, we decoded a multi-layered adaptive response characterized by synchronized immune evasion and profound metabolic reprogramming. Crucially, the full-length, unbiased nature of our transcriptomic module extends this regulatory landscape far beyond the coding genome, allowing us to capture critical non-coding elements that orchestrate this dormancy phenotype. For instance, we observed significant downregulation of the oncogenic long non-coding RNA (lncRNA) LINC00665^46^ which supports therapeutic efficacy, while LINC01644^47^ was markedly upregulated. This upregulation suggests a potential role for lncRNAs in mediating adaptive resistance, a layer of regulation that would remain invisible to proteome-only approaches.

However, this RNA-level regulation represents only a fraction of the cellular adaptive machinery. Beyond the transcriptome, our integrated dual-omics data provides a unique window into the “dark matter” of cellular regulation—the profound non-linear relationship between transcription and translation. Consistent with macroscopic proteogenomic studies, we observed a generally modest global correlation between mRNA and protein abundances in single cells. Rather than viewing this discordance as technical noise, we posit that it represents biologically vital post-transcriptional buffering mechanism. For instance, while some pathways, such as the complement system, exhibited concerted downregulation across both modalities, metabolic and DNA repair pathways demonstrated stark divergence. This indicates that under the acute therapeutic stress of chemotherapy, CSF-CTCs deliberately decouple protein stability from their transcriptional output, maintaining essential enzymatic pools to ensure survival despite a genome-wide transcriptional shutdown. These findings reinforce the paradigm that scRNA-seq alone is fundamentally insufficient to predict cellular phenotypes, particularly in highly dynamic clinical contexts where rapid proteostasis remodeling drives drug resistance.

While our integrated approach provides unprecedented insights, we recognize that capturing the ultra-low abundance regulatory proteome (e.g., transcription factors) remains a universal challenge in single-cell proteomics. Continued optimization of reaction miniaturization and automated LC-MS workflows will be essential to further scale analytical throughput and sensitivity. Regarding the cohort size, we recognize that the extreme clinical rarity and ethical constraints of obtaining serial CSF samples restricted our longitudinal analysis to a small number of patients. However, the robustness of our core findings, specifically the adhesive and metabolic reprogramming signatures was substantiated by their significant prognostic value in the large-scale TCGA-LUAD cohort, supporting their generalizability beyond our immediate sample set. Ultimately, the therapeutic targets and post-transcriptional regulators identified here serve as a robust proof-of-concept. Translating these vulnerabilities into clinical interventions will require a synergistic validation strategy that integrates experimental perturbations (e.g., CRISPR/Cas9 in patient-derived organoids) with AI-driven computational modeling, such as virtual cell knockout^48^. This hybrid strategy will allow for the rapid prioritization of high-confidence targets governing the malignant dormancy switch.

In conclusion, the integrated CLEAP-scMAPS pipeline stands as a robust, scalable, and device-free solution that bridges the critical gap between scarce clinical specimen acquisition and deep molecular phenotyping. By successfully resolving the stoichiometric landscape of rare CSF-CTCs, this platform paves the way for a new era of unbiased single-cell multi-omics. Future iterations of scMAPS could integrate isobaric labeling (e.g., TMT^49^) or faster LC gradients to further scale analytical throughput. Moreover, as demonstrated by our microneedle-based deterministic sampling of tissue micro-regions, scMAPS is inherently compatible with spatial biology. We envision a highly versatile framework, Spatial-scMAPS, that maps deep proteogenomic landscapes to their native architectural contexts, ultimately accelerating the discovery of precision therapies for devastating metastatic diseases.

## Methods

### Cell culture and collection

The hESC, L929, MCEC, BEAS and HEK293T cells were grown in a medium compatible with each cell line and all cultured for 5 days in a 37 °C incubator with 5% CO_2_. Then cells were detached with trypsin treatment after the supernatant was removed. Perform cell aggregate dissociation with strong pipetting. Cells were then rinsed twice with ice-cold PBS, pelleted by centrifugation at 1000 rpm for 5 min at 4 °C to remove the supernatant containing excess trypsin and culture media thoroughly. After that the cell pellets were resuspended in PBS for further use.

### Enrichment of CTCs in the cerebrospinal fluid of patients with lung cancer brain metastasis

All patients with lung cancer brain metastases have been fitted with Ommaya reservoirs.

Half of them received intracranial drug administration, while the other half had not yet received intracranial drug administration. The clinical CSF were collected from patients and infused into the slanted spiral polymer microfluidic chip to achieve label-free enrichment of CTCs. The study was approved by the Affiliated Nanjing Drum Tower Hospital of Nanjing University Medical School (Approval No: 2026-0189-01), and all patients provided written informed consent prior to participation. All experiments were performed in compliance with the Chinese laws and following the institutional guidelines.

### hESC cell lysates

For protein extraction, the above treated hESC cells were lysed in 1 mL lysis buffer (Solarbio, China) containing 10 mM EDTA, 150 mM NaCl, 1% Triton X-100, 1% phosphatase inhibitor cocktail and 100 x PMSF. Cells were sonicated in an ice bath for 10 min and repeated three times. Then the cell lysates were centrifuged at 12,000 rpm at 4 °C for 20 min and the supernatant was collected and transformed into a new EP tube. Protein concentrations were estimated using the BCA protein assay (Thermo Fisher Scientific).

For RNA extraction, the standard protocols were described previously^50^ and consists of sequentially adding TRIzol reagent, chloroform, isopropanol and 75% ethanol. Finally, 100 µL of ddH_2_O was added to dissolve the white substance in the tube, followed by mixing evenly with pipette, and stored at −80 ℃ until use.

### Capillary microneedle-based sampling platform of single cell

Prior to the cell capture operation, the 0.01% DDM pretreated PCR tubes were prepared by the addition of 2 μL of lysis buffer (0.5 μM Bio-dT_25_ probes (5’-Bio-CAAGGTTTTTTTTTTTTTTTTTTTTTTTTTVN-3’), 0.1% of DDM in 10 mM Tris, pH 8). The home-built capillary microneedle-based sampling platform originated from our group was used for single-cell picking up. The platform consisted of four parts including a x-y-z translation stage for position control, a system control unit, a microscopic camera system for cell observation and monitoring of the sampling process, and a tapered glass capillary (10 cm length, 200 µm i.d., 360 µm o.d., tip size, 30 µm o.d. (40 µm o.d. for CTCs)) for cell capturing operation. The target cell including cultured cell and CTCs was sucked into the microneedle under capillary force and connected to the compatible syringe a capillary tube to immediately dispense the cell into PCR tube. Finally, the isolated cells were immediately centrifuged at 1000 rpm for 1 min at 4°C to ensure them at the bottom of the tube and frozen at −80°C until use.

### Procedures of the scMAPS workflow

Isolated single-cells were lysed by the combination of freeze−thaw cycles and lysis buffer, which the samples were firstly frozen at −80°C for 10 min and thawed at 4°C for 3 min, then incubated at 56°C for 10∼15 min. Following lysis, the pre-added Bio-dT_25_ probes hybridized the mRNA molecules in the lysate with an incubation condition of 72°C for 3 min and 30°C for 10 min. Then, 1 µL streptavidin-microbeads (SA-beads, 10 µg/µL) were mixed with cell lysate and incubated at room temperature for 10 min with shaking at 800 ×rpm to enrich the mRNA/Bio-dT_25_ heteroduplex. The sample was centrifuged at 500 ×rpm for 3 s and placed on a magnetic rack for SA-beads enrichment, where cell lysate containing the protein supernatant was transferred into C18 filter 10 µL-tip for proteomics process, and the mRNA/Bio-dT_25_ heteroduplex bound SA-beads were used for subsequent transcriptomic analysis. Before use, rinse and activate the C18-tip by adding 20 µL of 80% ACN or MeOH, then centrifuge at 100 ×g for 1 min to ensure the C18 pads are secured and stabilized.

For single-cell transcriptomic analysis, the SA-beads bound with mRNA/Bio-dT_25_ heteroduplex were washed twice with 0.2 × SSC, resuspended with 2.5 µL of ddH_2_O and incubated at 80°C for 2 min to elute the mRNA. Then, the sample was instantaneously centrifuged and immediately placed on a magnetic rack for SA-beads enrichment. The supernatant containing mRNA was then transferred to a new PCR tube preloaded with 0.5 μl of 10 μM CBTi-oligo primer, incubated at 72°C for 3 min and directly placed back on ice. The system was then supplied with 10 µL of one-step RT-PCR reaction mix containing 1.2 µL dNTP (10 mM), 0.252 µL RNase Inhibitor (40 U/µL), 0.1 µL DTT (0.1 M), 0.328 µL MgCl_2_ (279 mM), 0.18 µL dCTP (100 mM), 2 µL Betaine (5 M), 0.368 µL CBTi-TSO (50 µM), 0.172 µL of Maxima H Minus (200 U/µL, Thermo Scientific) and 5 µL of KAPA HiFi HotStart Ready Mix (2×, Roche). The following RT-PCR program was used: 60 min at 50 ℃, 98 ℃ for 3 min, then 20-22 cycles (98 ℃ for 20 s, 67 ℃ for 20 s, and 72 ℃ for 6 min) and 4°C hold on the thermocycler.

Then, 1 ng or 5 ng unpurified cDNA working dilution was transferred to a new microtube and added into tagmentation reaction mix which contained 4 µl Tagment buffer (5×), 0.5 µl Barcode A_m_ Tn5 (40 ng/µl), 0.5 µl Barcode B_n_ Tn5 (40 ng/µl) for 1 ng cDNA (or 1.25 µl Barcode A_m_ Tn5 (40 ng/µl), 1.25 µl Barcode B_n_ Tn5 (40 ng/µl) for 5 ng cDNA), and nuclease-free water to 20 µl final volume. Next, each sample was tagmented, multiplexed collection, and library amplification based on the previously reported CBTi-seq multiplexed transcriptome sequencing technology^20^.

For single-cell proteomics process, the proteins in tip were digested by adding 1 µL of 10 ng/µL Pierce™ Trypsin/Lys-C Protease Mix (Thermo Scientific) in 50 mM ABC and incubating at 37°C for 5 h. After digestion, a final concentration of 1% FA was added to stop the digestion. The digested peptides were centrifuged on the spintip device (800 g for 60 s) and eluted with 6 µL 98% ACN/0.1 FA buffer (twice) into glass insert tube. Finally, these peptide solutions were dried with a SpeedVac and reconstituted using 7 μl of 0.1% FA for LC-MS/MS analysis.

### LC-MS/MS analysis

The DDA samples were separated at a flow rate of 150 nL/min and analyzed with a linear 130-min gradient of 8-30% mobile phase B (0.1% FA in ACN) using an UltiMate 3000 RSLCnano LC system and Orbitrap Eclipse Tribrid MS (Thermo Scientific, Xcalibur Ver. 4.3.73.11). Electrospray voltage of 2.2 kV was applied at the ionization source to generate electrospray and ionize peptides. The ion transfer capillary was heated at 300°C for droplets desolvation. Data were acquired with full MS scans from m/z 375-1500 at a resolution of 120,000 (at m/z 200) and automatic gain control (AGC) target of 1E6. Precursor ions with charges of +2 to +7 were isolated with an isolation window of 4, an AGC target of 1E5, a maximum injection time of 300 ms, and fragmented by higher energy collisional dissociation (HCD) of 30% for MS/MS analysis. A dynamic exclusion with duration of 30 s was employed to minimize repeated sequencing, and mass tolerance of ±10 ppm was used. For DIA, the peptide samples were separated on a Vanquish Neo UHPLC system (Thermo Fisher Scientific) with a 30 min gradient and analyzed via an Orbitrap Astral mass spectrometer (Thermo Fisher Scientific), following a previously established protocol^21^.

### Database searching and data analysis

The raw transcriptome sequencing data was subjected to quality assessment by FastQC, barcode sequence swapping, and trimmed to generate clean data, respectively. The reads were then aligned to the human genome (GRCh38) or mouse genome (GRCm38) using STAR software (version 2.7.11b). After alignment, the DigitalExpression function in Drop-seq_tools-3.0.2-0 was employed with specified filtering conditions to select the digital expression matrix corresponding to the specified Barcode. Finally, the numbers of total reads were summarized to normalize the gene expression. The minimum of 1 TPM reads at least 50% samples were used to filter genes with low abundance for all cluster experiments with BEAS, HEK293T, and CSF-CTCs cells. In cases where complete data was needed, imputation was performed with k-nearest neighbors. Differentially expressed genes were analyzed using DESeq2 software, where genes that met the thresholds of adj.p-value < 0.05 were defined as DEGs. Coverage across the gene body was calculated by RseQC. Reads Coverage and Sashimi plot of sequencing data aligned at the POU5F1 and Col1a1 were visualized using the IGV (version 2.17.4) genome browser.

Protein LFQ identification from the DDA dataset was searched by Proteome Discoverer Software (version 2.4) with the setting against the UniProt proteome database (Homo Sapiens: 20,443 entries; Mus musculus: 17,201 entries). The parameters of the database search are as follows: 2 missed cleavage sites, mass tolerances for precursor ions and fragment ions of 20 ppm and 0.02 Da, respectively. The methionine oxidation and protein N-termini acetylation were set as variable modifications. For DIA data, the MS raw files were analyzed with DIA-NN software (version 1.9.2) in library-free search mode using default settings. A false discovery rate (FDR) of 0.01 was filtered to reduce false matching. Based on the results obtained, Perseus software (version 1.5.5.3) was used to perform quantified data analysis and extraction. The LFQ protein abundances were log-transformed and filtered to contain at least 70% valid values in each group for further test. In cases where complete data was needed, the missing values were imputed by normal distribution in each column. The statistical analyses were all processed and visualized with Origin (version 2021), GraphPad Prism (version 8.0.2), and R studio (version 4.3.2). The PPI networks were analyzed by STRING database and visualized by Cytoscape software (version 3.9.1).

## Acknowledgments

This work was supported by the National Natural Science Foundation of China (82361138570, 81827901) and National Natural Science Foundation of China Young Scientists Fund (No. 82304567). We acknowledge the Analysis and Testing Center of Southeast University and Tiannan Guo group of Westlake University for MS instrumentation and technical support. Fig. 1 and Fig. 3a was created with BioRender.com.

