## Supplementary information for "Unbiased Single-Cell Transcriptome-Proteome Co-Profiling Reveals Malignant Dormancy and Post-Transcriptional Buffering of CTCs"

**Methods**

**Carboxylated oligo(dT)-beads preparation.** The carboxylated magnetic microbeads used in all experiments were commercially available (KBsphere MagCOOH). The microbeads have an average diameter of 1 µm and were coated with a hydrophilic surface. For oligo-dT capture probe assembly, 6.25 µL of microbeads (50 mg/mL) were added to the 0.2 mL PCR tubes, rinsed twice with 0.1 M MES, and further resuspended in 50 µL of MES (0.1 M). Then 6.25 µL of EDC (50 mg/mL) and 5 µL of 100 µM oligo-dT capture probe (5′-NH2 C6-CAAGGTTTTTTTTTTTTTTTTTTTTTTTTTVN-3′) were added to the PCR tubes and incubated at 28 °C for 4 h with 1500 rpm to achieve the effective assembly of oligo-dT. After that the supernatant solution was removed by magnetic separation and the oligo-dT microbeads were then washed three times with 100 µL of TET buffer containing 2 µM Tris-HCl, 5 µM EDTA, 0.1% Tween-20, and RNase-free distilled water, then resuspended in 31.25 µL of TET buffer. Finally, the synthesized oligo-dT microbeads were incubated with 2% BSA on a rotator for 4 h at 20 °C to prevent non-specific absorption sites, and then the BSA solution was exchanged with 31.25 µL of 1 ×PBS buffer prior to experiment.

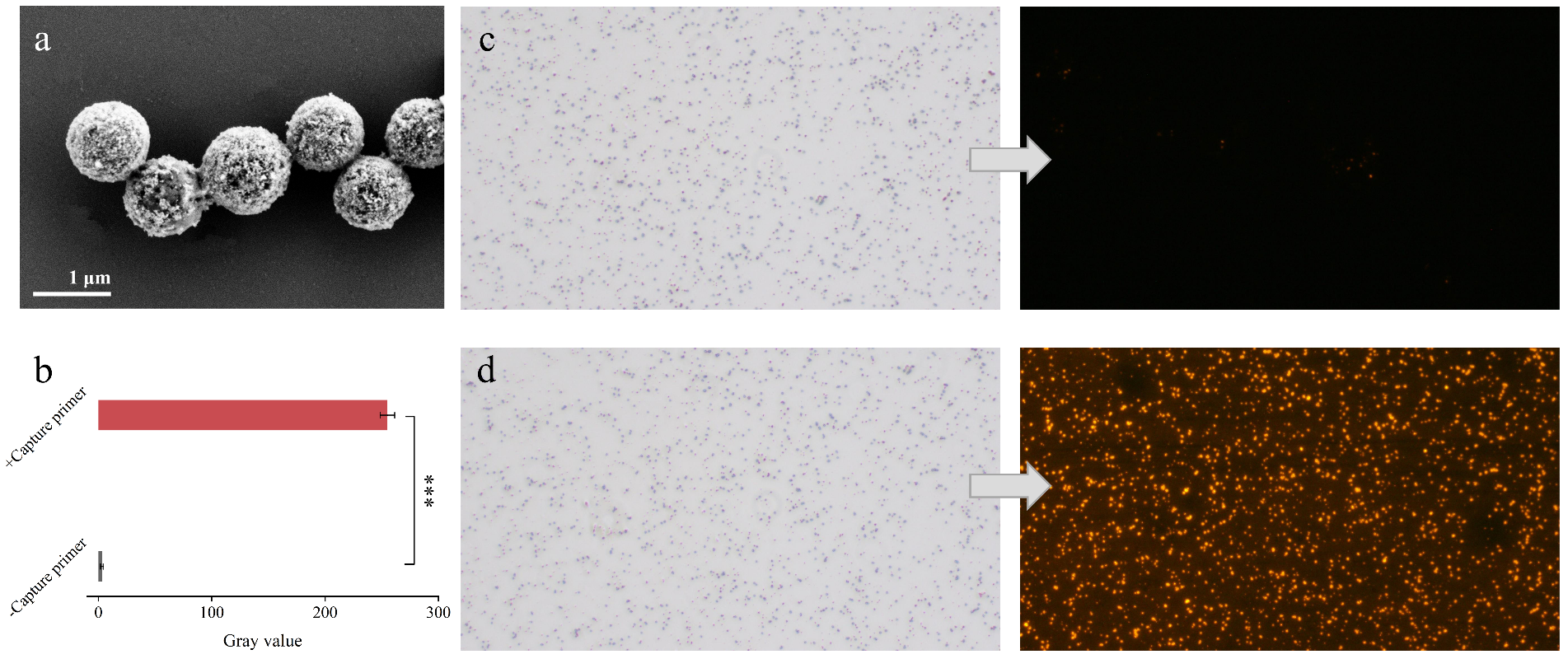

**Fig. 1. Image characterization of the SA-beads enrichment capability.** a, SEM image of SA-beads. b, The gray values through ImageJ software. The bright/fluorescence image of bare SA-beads (c) and poly-dA probe/Biotin-dT_25_ heteroduplex immobilized SA-beads (d).

Scanning electron microscopy (SEM) revealed that the SA-beads possess a uniform spherical structure with a mean diameter of 1 μm (Fig. 1a). To verify hybridization between the biotin-dT_25_ probe and poly(A^+^) RNA on the SA-beads, we incubated the beads with a fluorescent poly-dA probe modified with Cy3 at the 5’ end and analyzed them via fluorescence microscopy. We observed that the fluorescence intensity of the SA-beads immobilized with poly-dA/biotin-dT_25_ heteroduplex was significantly higher than that of bare SA-beads (Fig. 1c, d), and the significant difference was further confirmed by quantifying gray values through ImageJ software (Fig. 1b), demonstrating the high capture efficiency of the biotin-dT_25_ functionalized SA-beads and their suitability for robust downstream molecular fractionation.

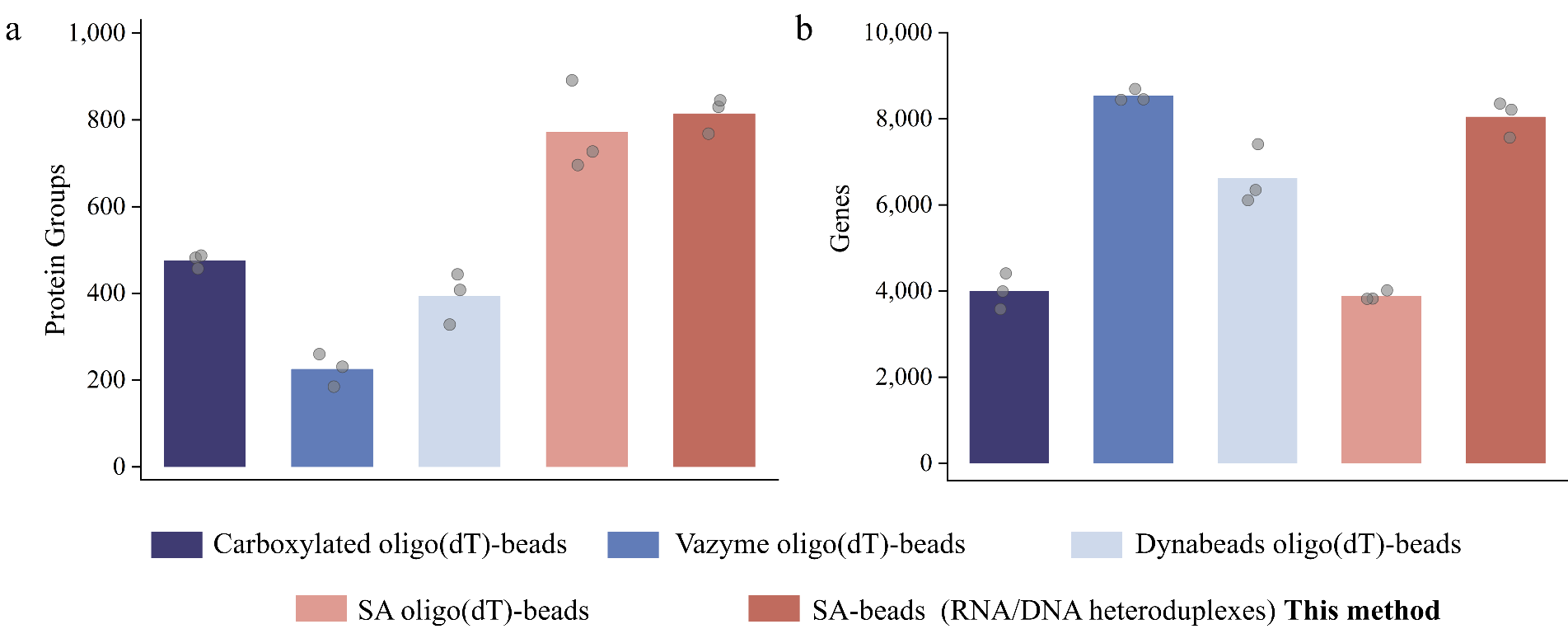

**Fig. 2. The impact of different types of magnetic beads and their enrichment efficiency on multi-omics analysis.** Protein groups (a) and genes (b) identification under the 10 pg RNA and 1 ng proteins extracted from hESC cells.

The results indicated that commercial options, such as Vazyme and Dynabeads oligo(dT)-beads, substantially compromised protein group identification, which may be attributed to the non-specific adsorption of proteins caused by exposed sites on the bead surface. Conversely, the streptavidin (SA) protein layer on our SA-beads, combined with blocking treatments, effectively mitigated this issue (Fig. 2a). Notably, self- assembled carboxylated/SA oligo(dT)-beads detected only ~50% of genes recovered by our method, indicating suboptimal poly(A^+^) RNA capture efficiency at solid-liquid interface (Fig. 2b).

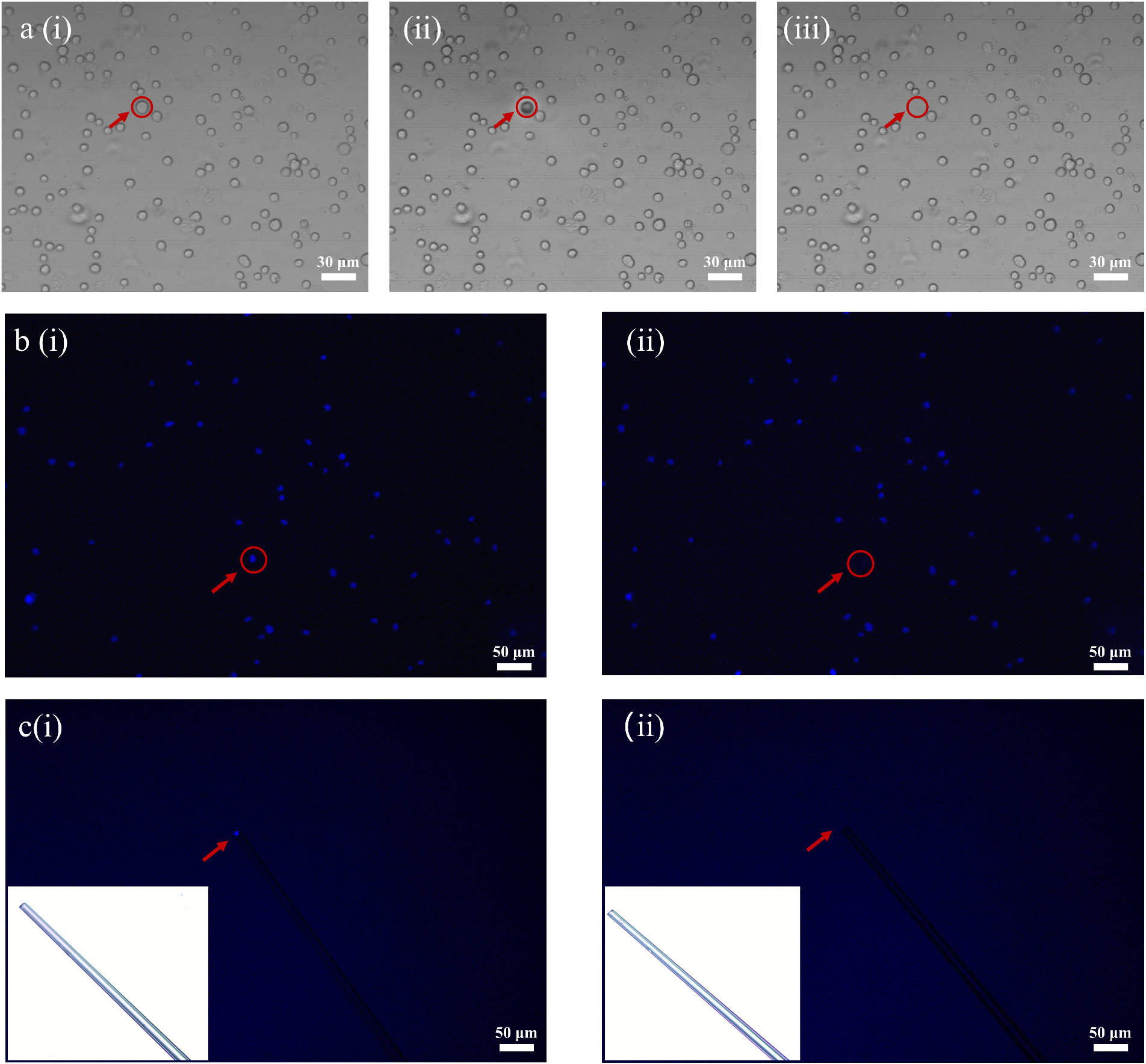

**Fig. 3. The characterization diagram of single-cell picking using capillary microneedle platform.** a, In the microscopic bright-field view, i-iii respectively represent the state of freely dispersed single cells before collection, when the microneedle is close to the needle tip, and after the microneedle collects the single cell. b, In the fluorescence microscopic view, i-ii respectively represents the fluorescence cells stained with DAPI before and after picking. c, In the fluorescence microscopic view, i represents the fluorescence characterization of the microneedle after collecting the cells, and ii represents the fluorescence characterization of the transferred cells after collection. Among them, the lower left corner of the Fig. c represents the characterization of the microneedle before and after cell transfer in the bright-field view.

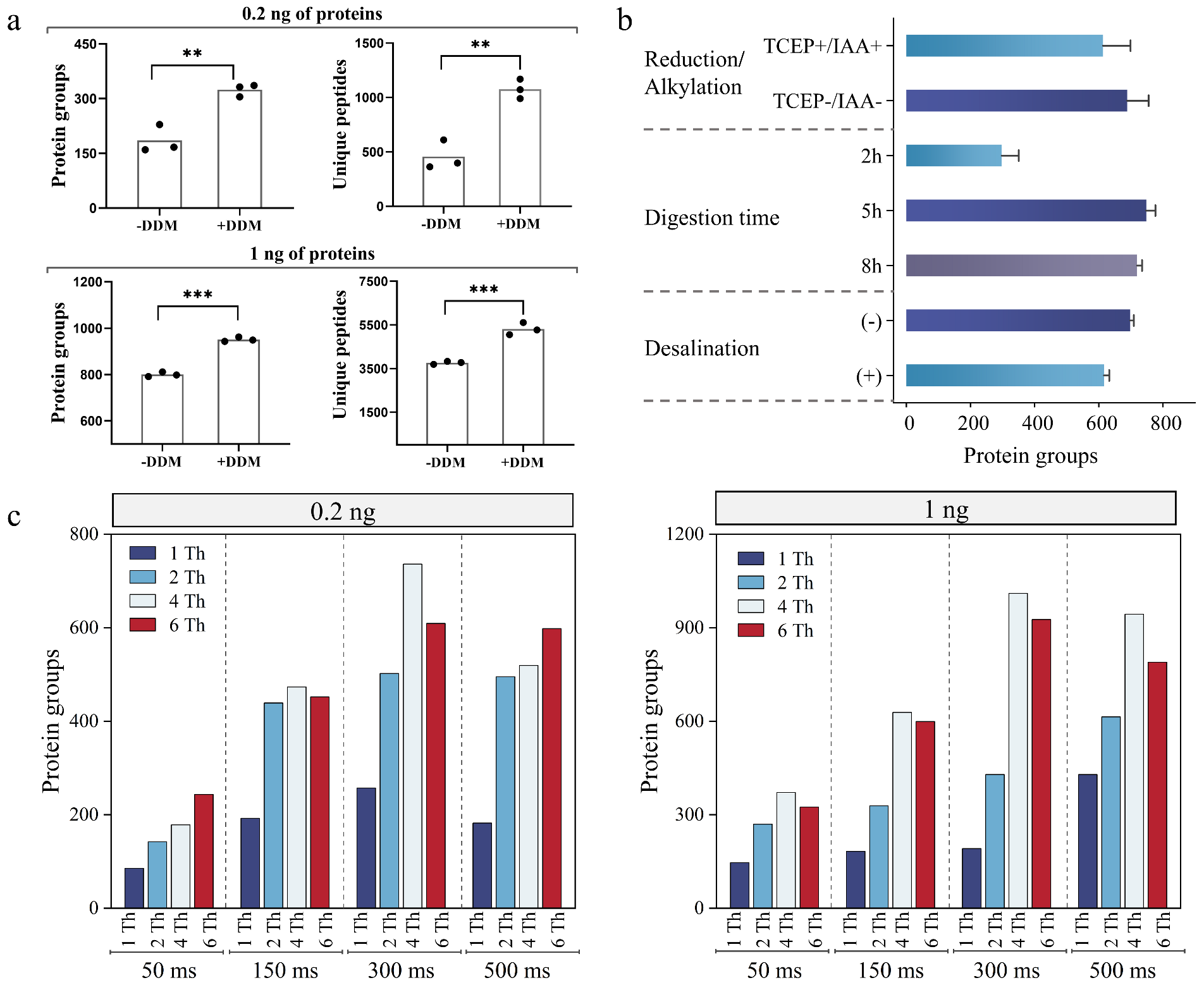

**Fig. 4. Optimization and performance of DDA-MS.** a, The impact of DDM-coated PCR microtube on the proteome identification using 0.2 ng and 1 ng of hESC proteins, respectively. b, The optimization of reduction/alkylation steps, digestion time (2-8 h), and desalination step. c, Optimal isolation window (IW) and maximum ion injection time (ITmax) using 0.2 ng and 1 ng of hESC proteins, respectively.

We first evaluated the impact of using DDM-coated PCR microtubes on proteome coverage. Compared to standard PCR microtube, DDM-coated PCR tubes yielded a remarkable increase in unique peptide and protein groups detections. Specifically, we observed increases of 135.7% and 75% for 0.2 ng protein inputs, and 40.9% and 18.9% for 1 ng inputs, respectively (Fig. 4a). The results indicate that coating with the nonionic surfactant DDM significantly mitigates protein loss by reducing hydrophobic adsorption to the contact surface.

Furthermore, streamlining the ultrasensitive proteomic sample preparation revealed unexpected optimizations (Fig. 4b). Contrary to bulk proteomics dogma, omitting reduction/alkylation steps (5 mM TCEP/15 mM IAA) increased protein identification, suggesting chemical derivatization impairs trace-level recovery. We next calibrated trypsin digestion duration, observing peak sensitivity at 5 h, with longer incubations (8 h) reducing yields due to peptide adsorption losses. In addition, eliminating desalting boosted recovery by 10% versus in-staged-tip methods, as salts were negligible in micro-volume samples. Collectively, this redefines minimalistic processing for proteome in the scMAPS workflow. Optimizing MS parameters proved equally critical for trace-level peptide detection, as insufficient precursor ions accumulation often precludes the generation of quality MS2 spectra. Here we explored two crucial factors for optimal precursor ions acquisition and peptide detection, i.e. isolation window (IW) and maximum ion injection time (ITmax) under the DDA mode. Expanding ITmax (50-300 ms) and IW (1-4 Th) can significantly improve the recognition rate, which is attributable to the enhanced ion transmission and accumulation. However, spectral complexity from reduced ion purity compromised recognition above 4-Th IW, highlighting the trade-off between sensitivity and specificity (Fig. 4c). Consequently, we established an ITmax of 300 ms and an IW of 4 Th as optimal parameters for trace proteomics, maximizing depth without sacrificing reliability.

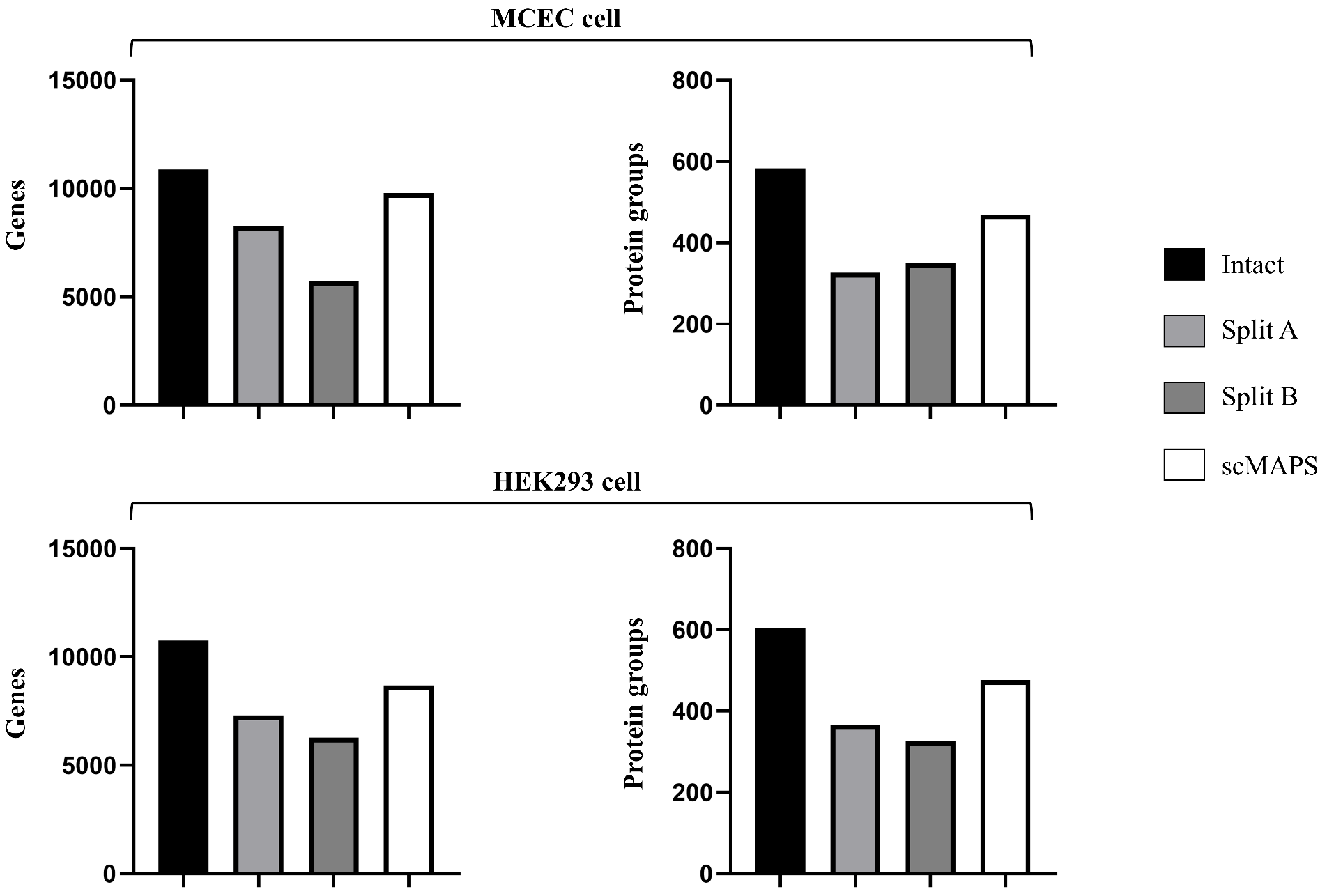

**Fig. 5. Comparison of the sample splitting technique and the scMAPS approach in terms of multi-omics identification capabilities from single MCEC and HEK293 cell analysis.**

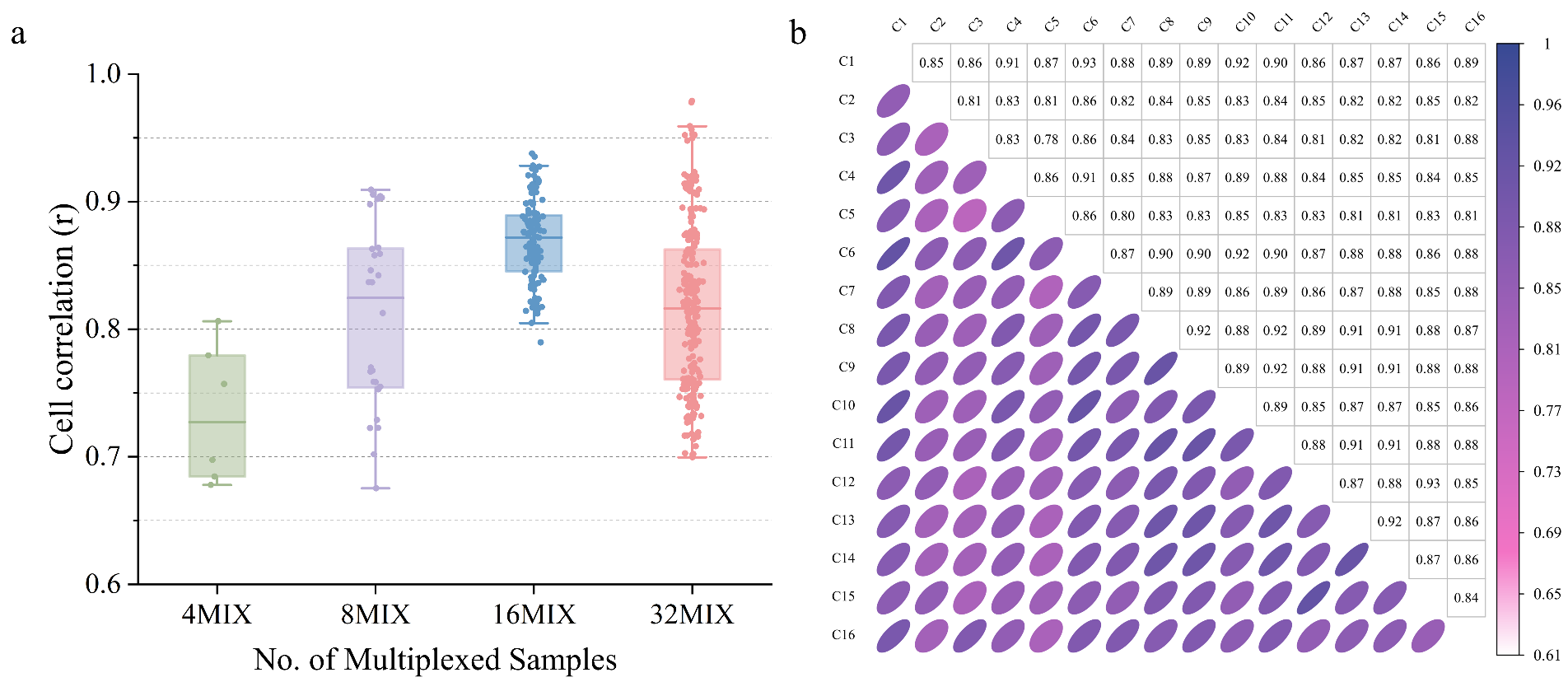

**Fig. 6. Inter-sample quantitative correlations by CBTi-seq.** a, inter-sample quantitative correlations under the conditions of different multiplexed hESC cells (4, 8, 16, and 32MIX). b, The quantitative correlations heatmap of 16MIX hESC cells.

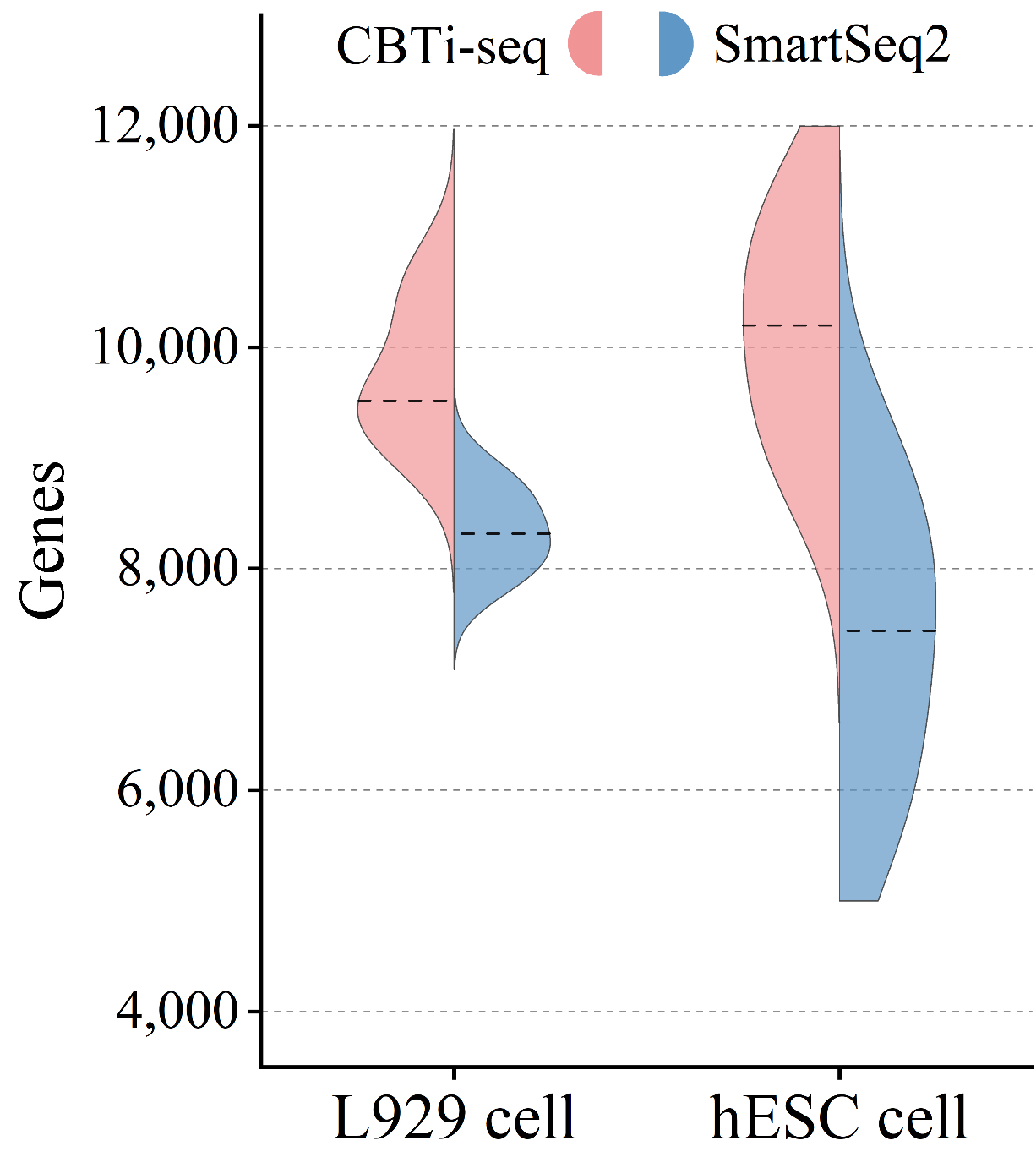

**Fig. 7. Comparison of sensitivity to detect genes in L929 cells and hESC cells processed with CBTi-seq and Smart-seq2 method (n=3 for each method).**

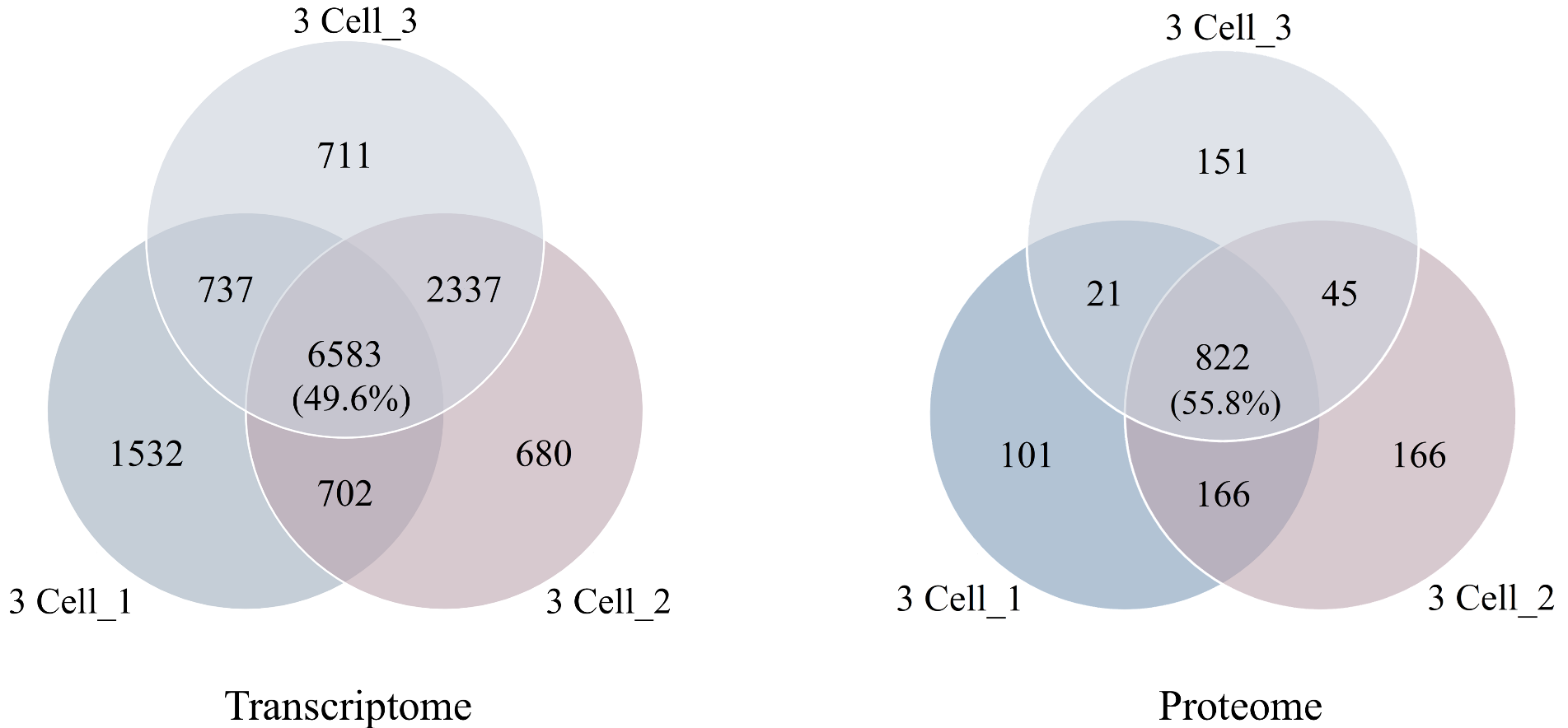

**Fig. 8. Venn diagram of the identified genes (left) and proteins (right) between the triplicate analysis of 3- HEK293T cells loadings.**

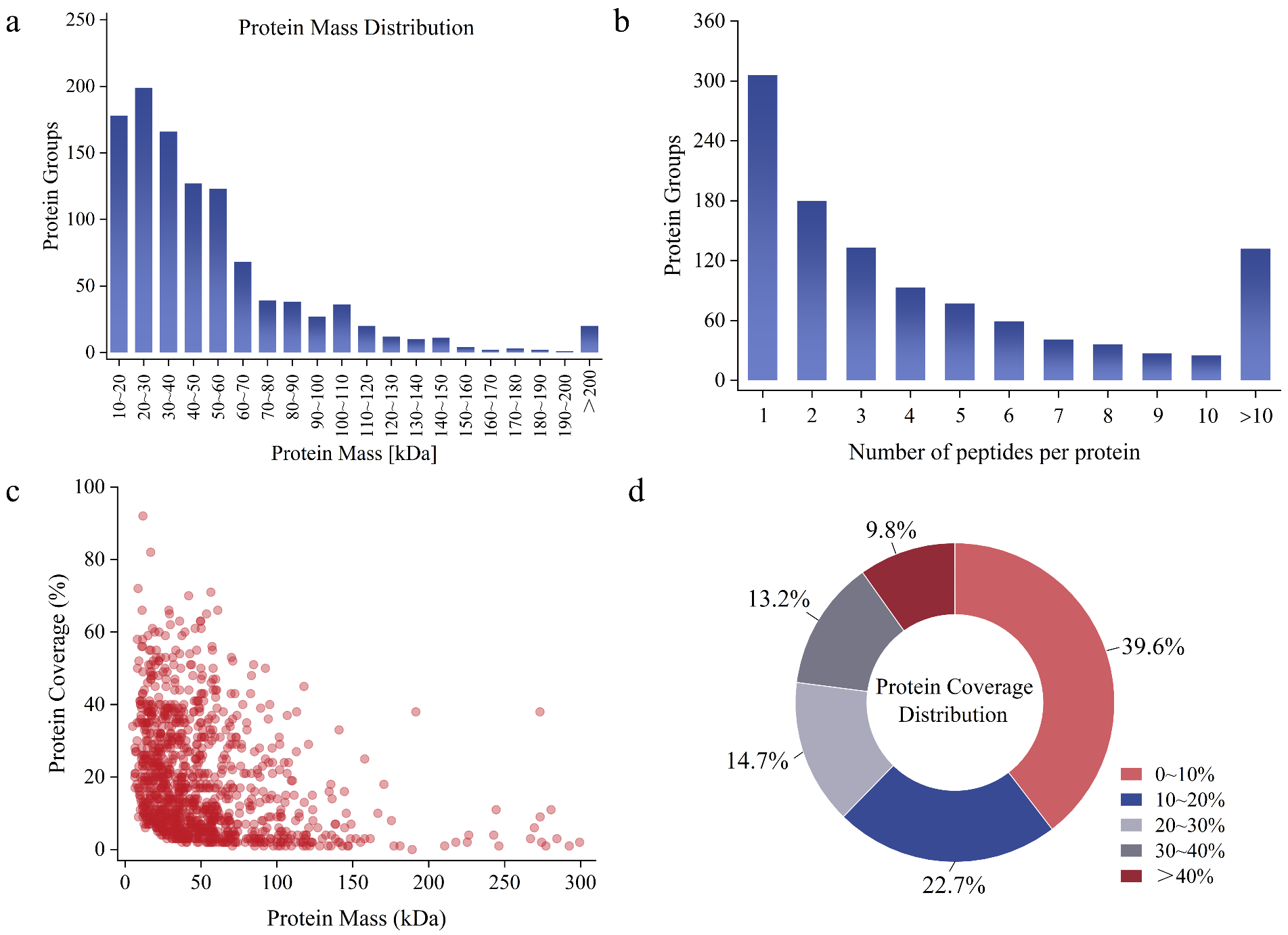

**Fig. 9. Quality control results of the DDA-MS in 3-HEK293 cell.** a, Protein Mass Distribution. b, Number of peptides per protein. c, Protein coverages of several protein Mass. d, Protein coverage distribution.

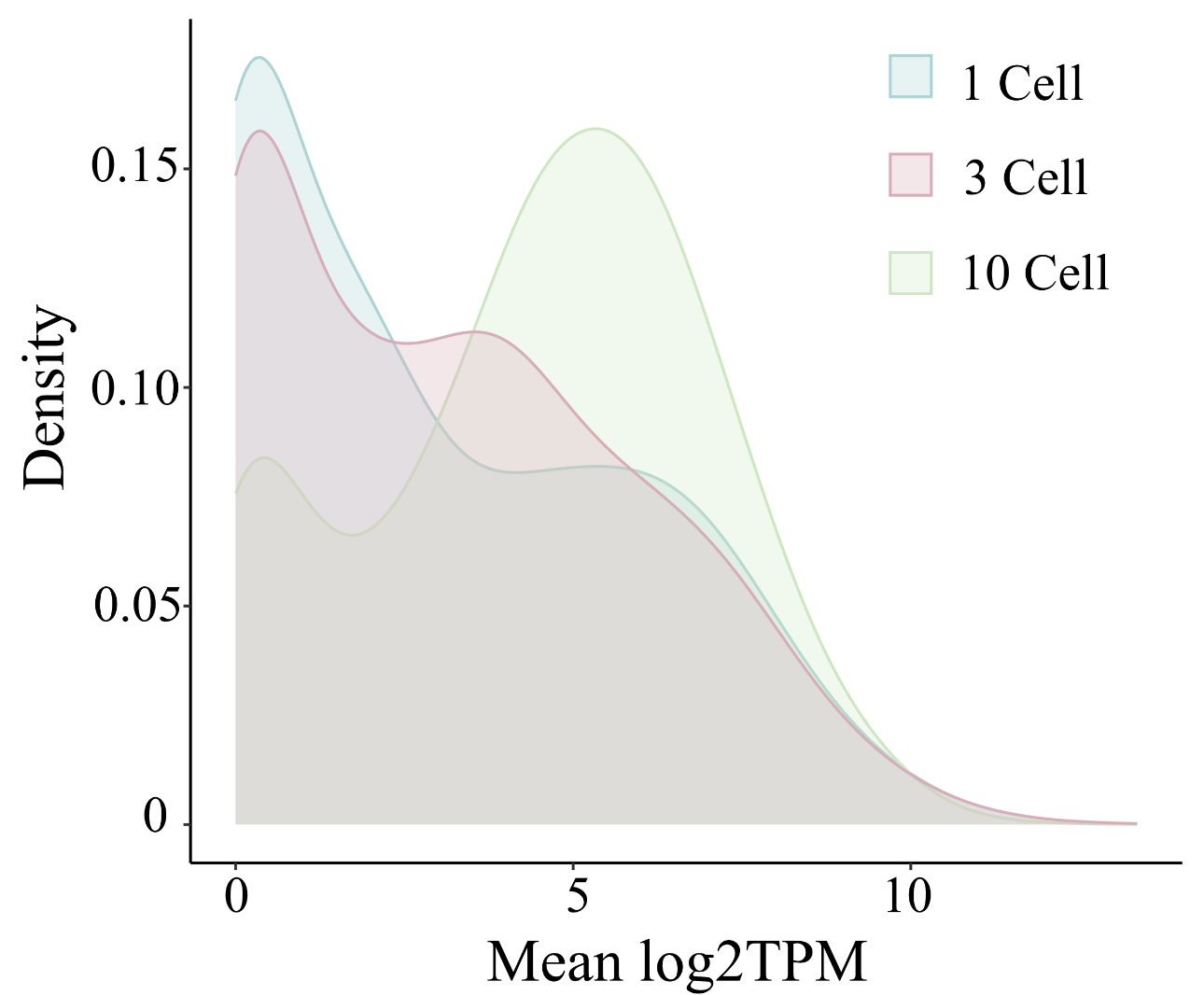

**Fig. 10. Density plot illustrated the distribution of mean log₂-transformed TPM (transcripts per million) values across different cell (1, 3, and 10 cell) quantities.**

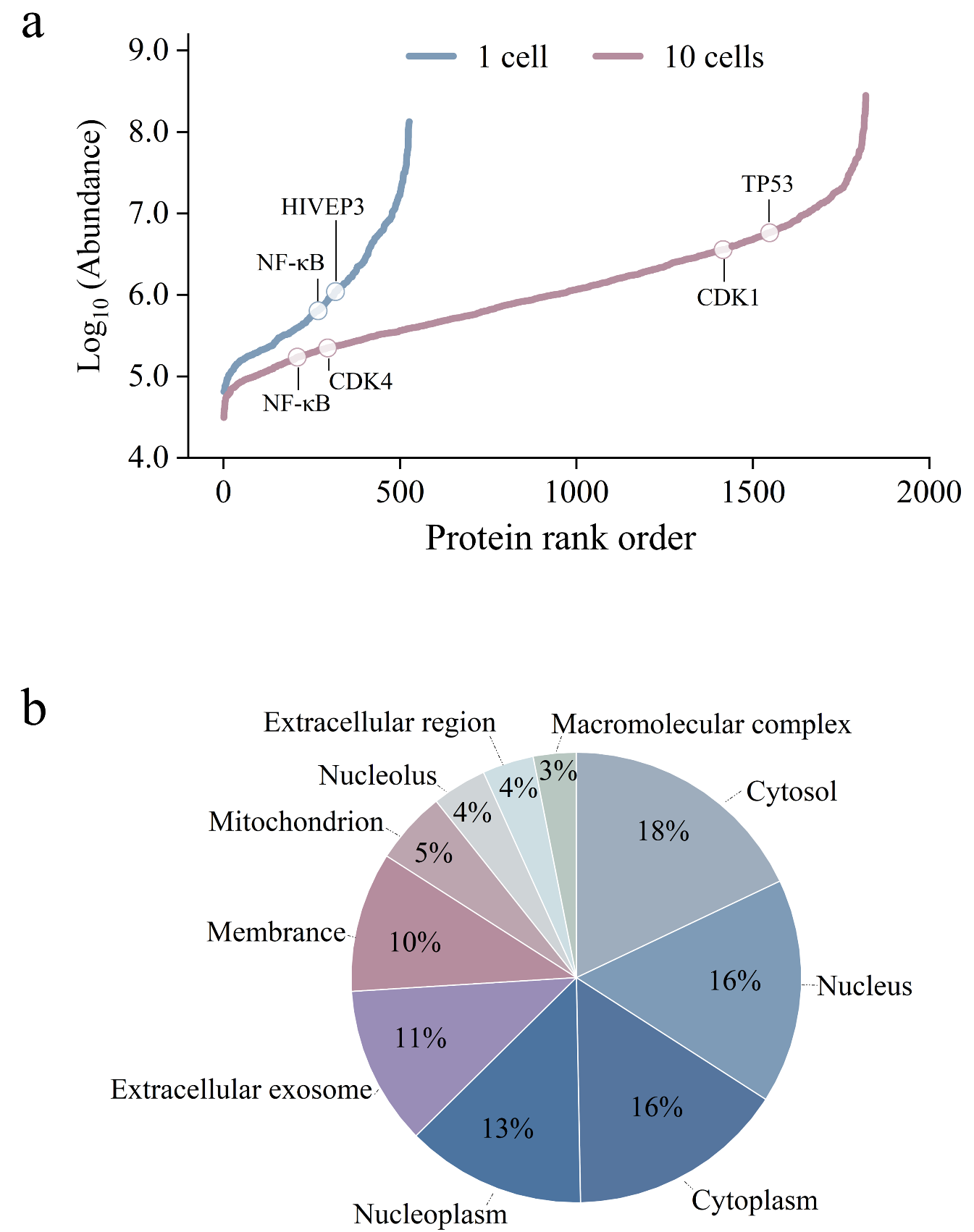

**Fig. 11.** a, Assessment of dynamic range based on protein abundance rank and annotation of selected typical proteins. b, GO-CC distributions of proteins identified in the 3-cell groups.

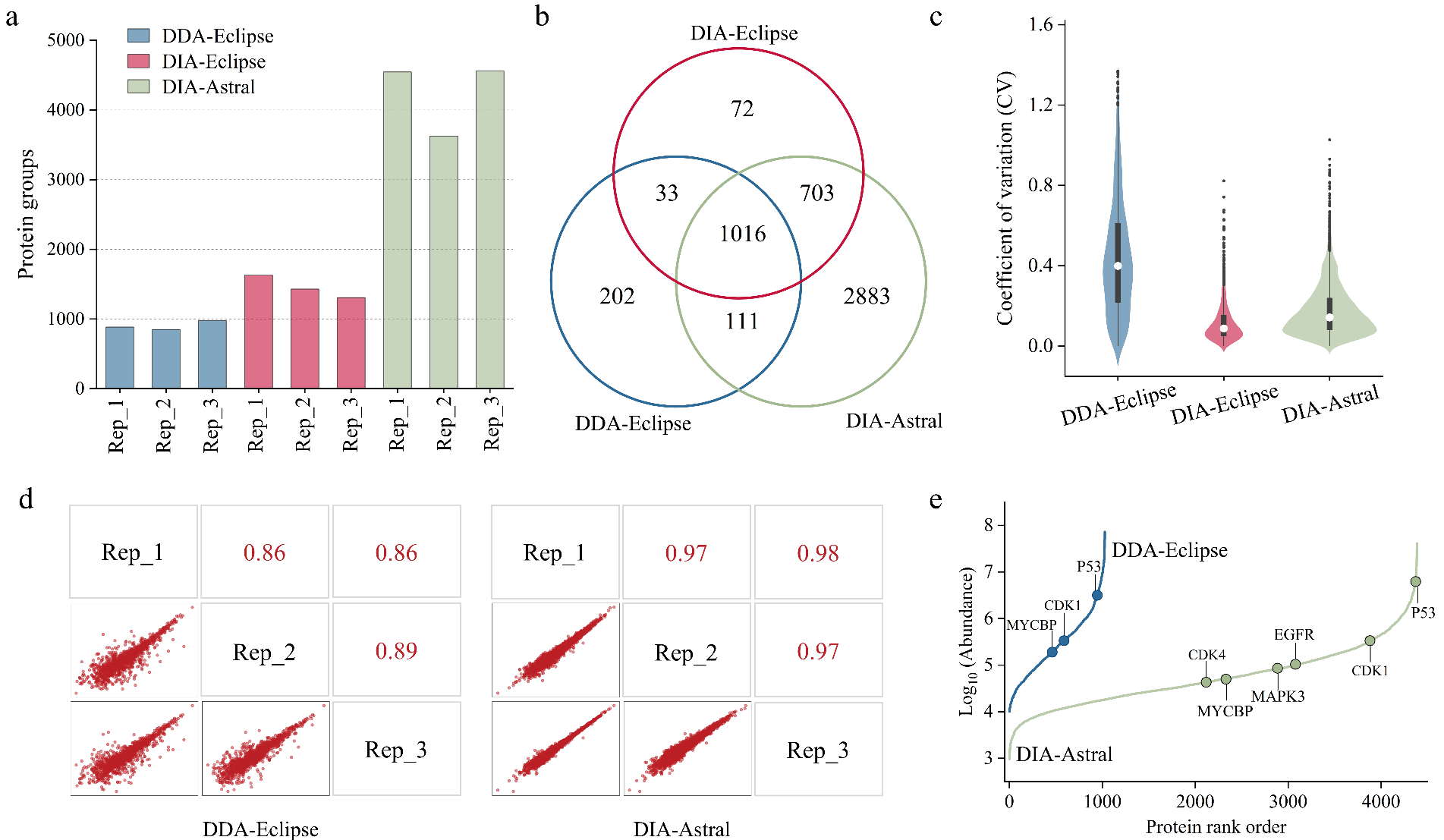

**Fig. 12. Test for the repeatability of 3 consecutive analysis of 250 pg standard HeLa peptides using the LC-MS system in DIA-Eclipse, DIA-Astral and DDA-Eclipse modes.** The results of protein identification number (a), overlap of protein identification (b), distributions of the coefficients of variation (c) and their Pearson correlation coefficients of pairwise analysis (d) are shown. e, Assessment of dynamic range based on protein abundance rank and annotation of selected proteins related to cancer.

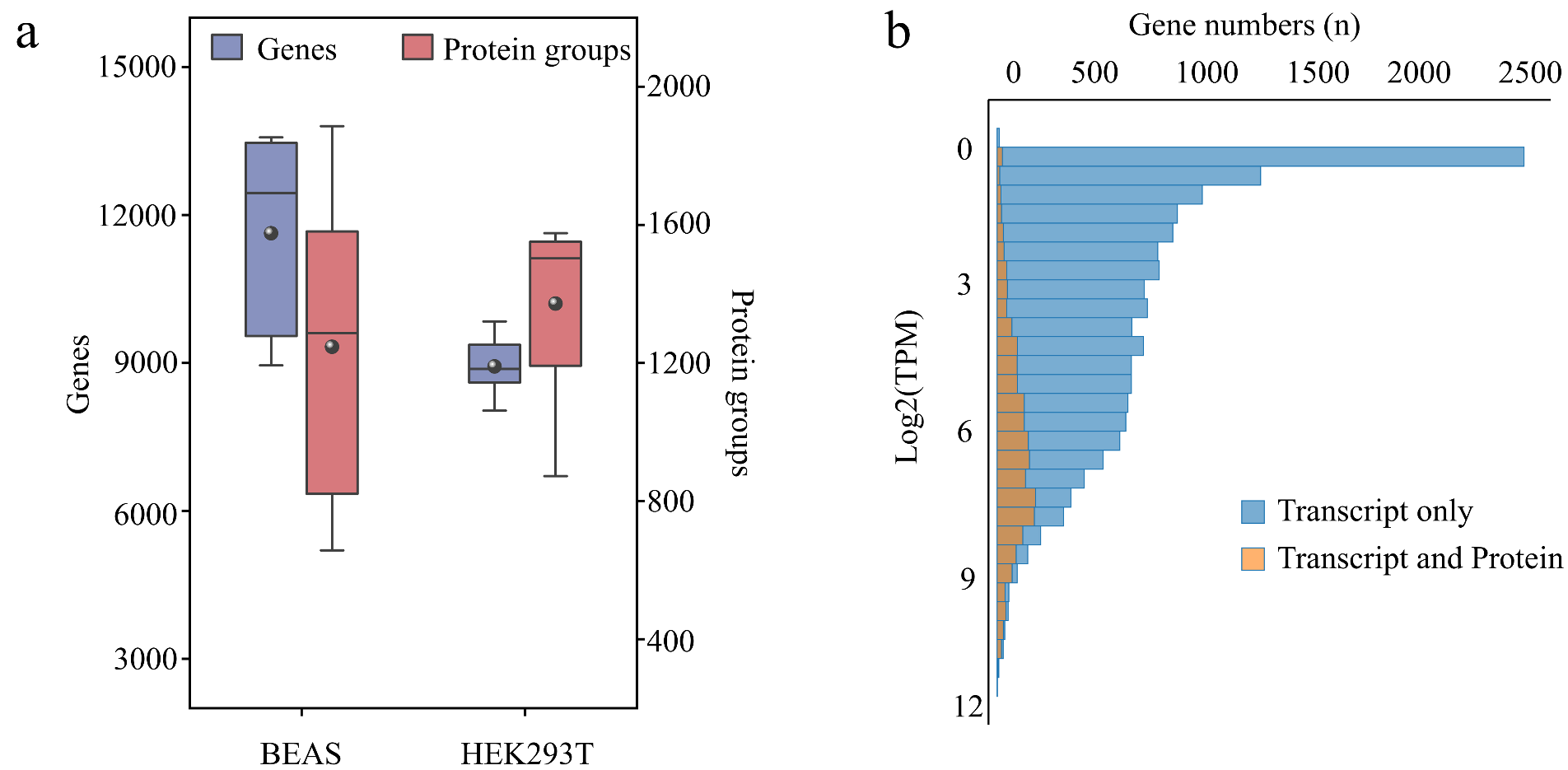

**Fig. 13.** a, The average number of detected genes and protein groups in single BEAS (n = 10) and HEK293T (n = 9) cells (protein groups are detected under the DIA mode). Box plots show the median (center), 25th/75th percentile (lower/upper hinges) and1.5× interquartile range (whiskers). b, Histogram of all genes detected with transcripts per million (TPM) > 0.1 at transcript and/or protein level, arranged by the average log2TPM of each gene detected.

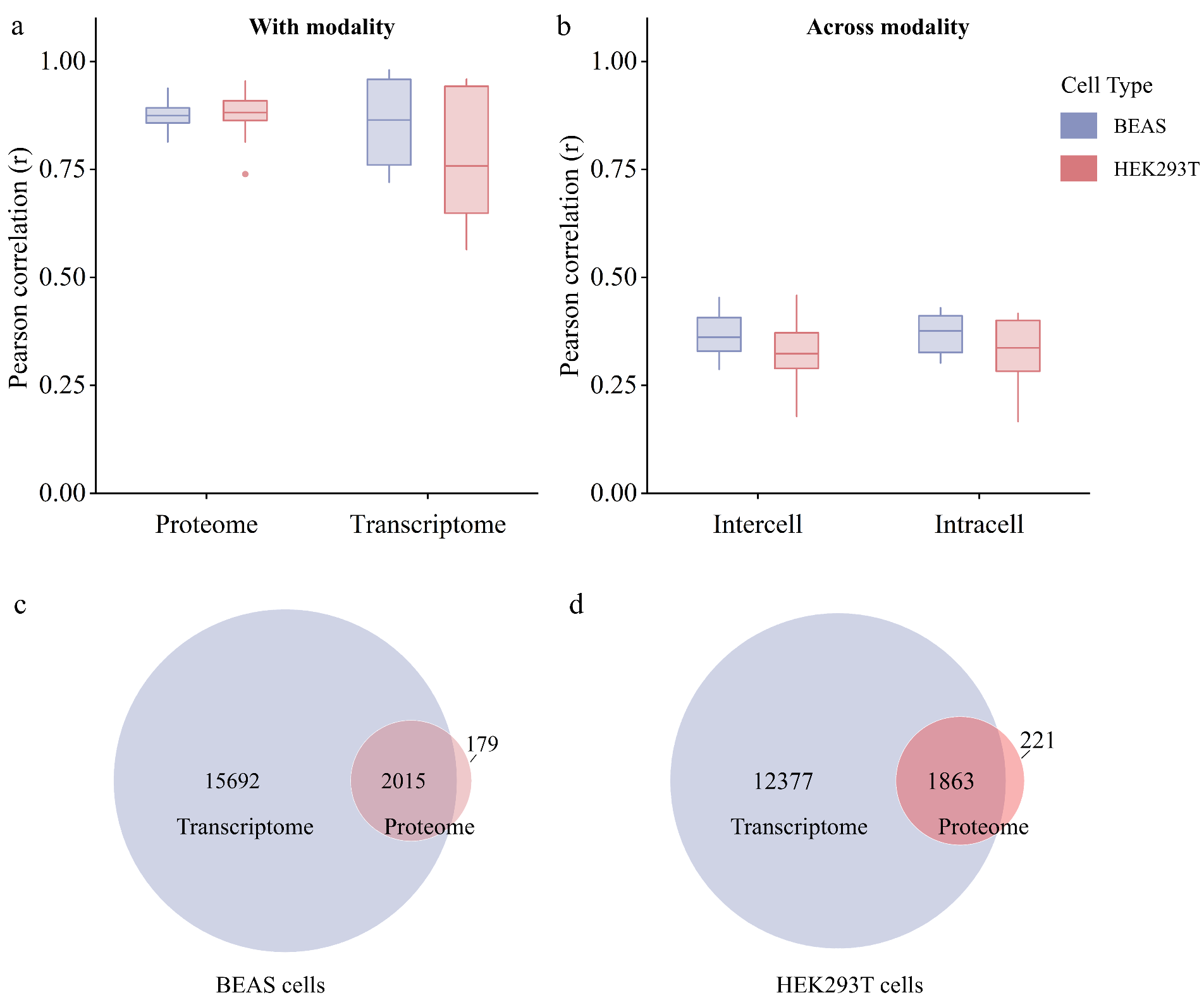

**Fig. 14. Distributions of correlations and overlapping identifications (BEAS and HEK293 cells).** Box plots showing the distributions of Pearson correlations, separated by cell types (n = 10 for BEAS and n = 9 for HEK293) within modalities (proteome and transcriptome) (a) and across modalities (b). (c) Overlap in gene and protein identifications from BEAS cells. (d) Overlap in gene and protein identifications from HEK293 cells. The center line in the boxplot represents the median while the box boundaries extend from the first (25%) and third (75%) quartile, representing the interquartile range (IQR). Boxplot whiskers extend to 1.5*IQR.

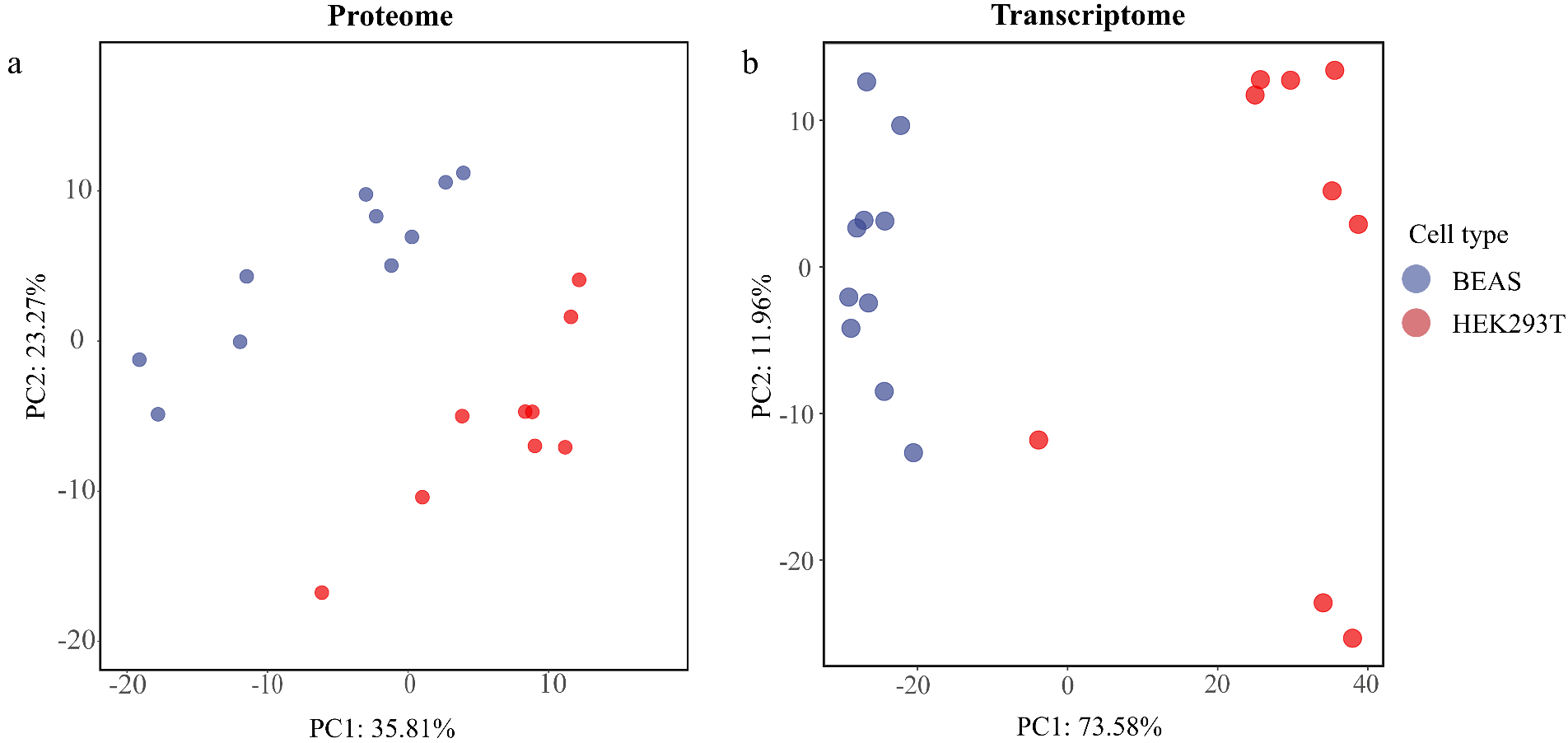

**Fig. 15. PCAs of BEAS and HEK293T cells from proteome and transcriptome.** (a) PCA of BEAS (n = 10) and HEK293T (n = 9) cells using proteome data and (b) transcriptome data.

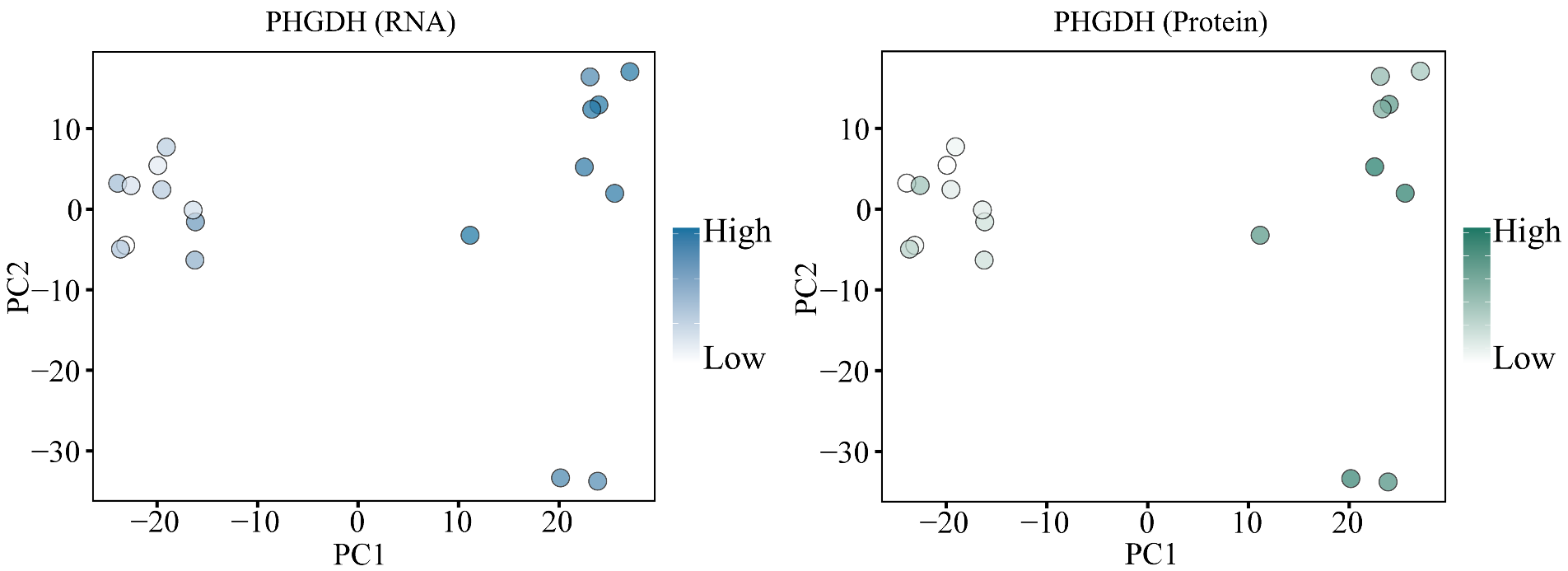

**Fig. 16. Visualization mapping of the gene (blue) and protein (green) abundance of PHGDH.**

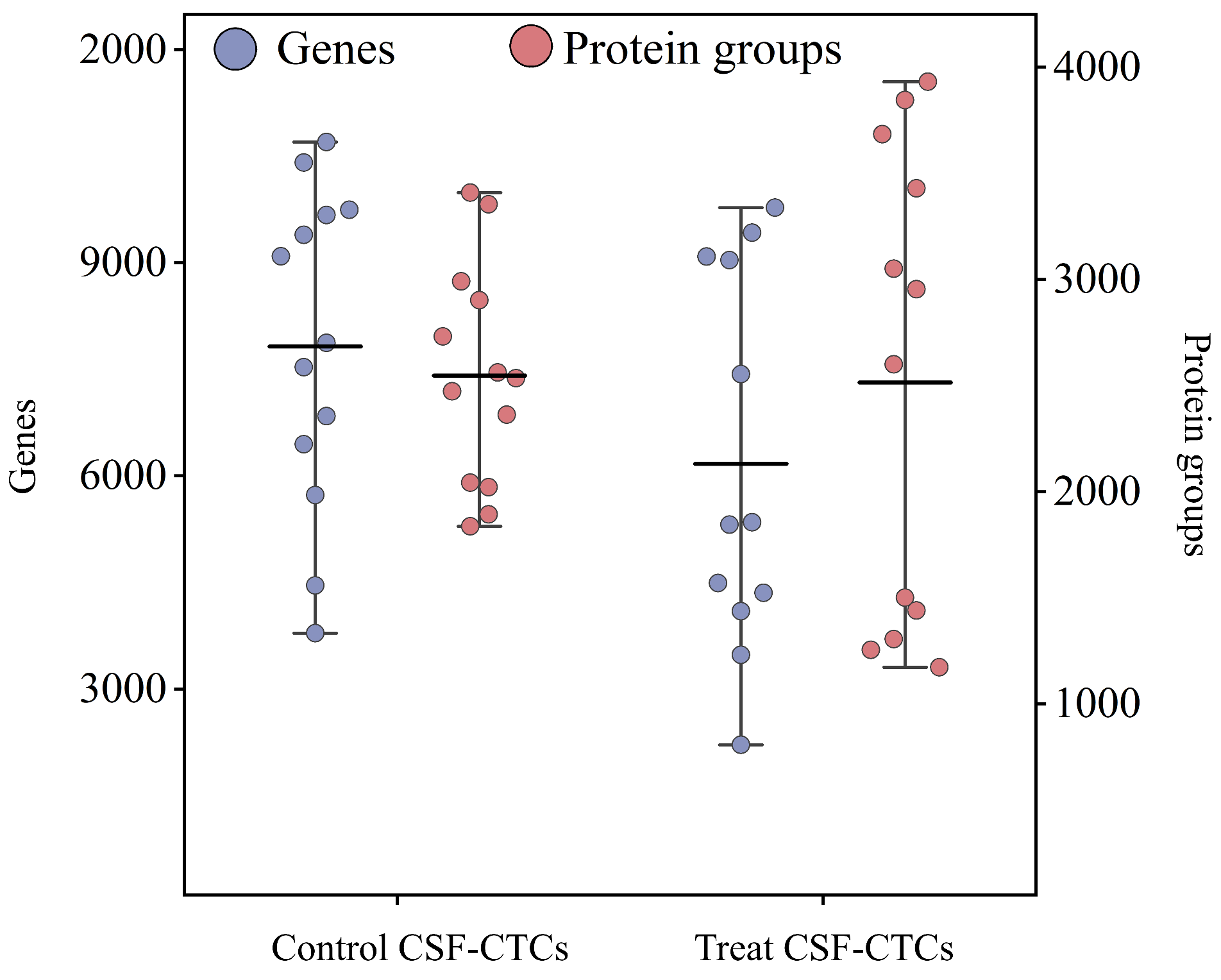

**Fig. 17. The number of genes and proteins identified in the control CSF-CTCs group and the treat CSF-CTCs group.** The horizontal line represents the average value. Boxplot whiskers extend to 1.5*IQR.

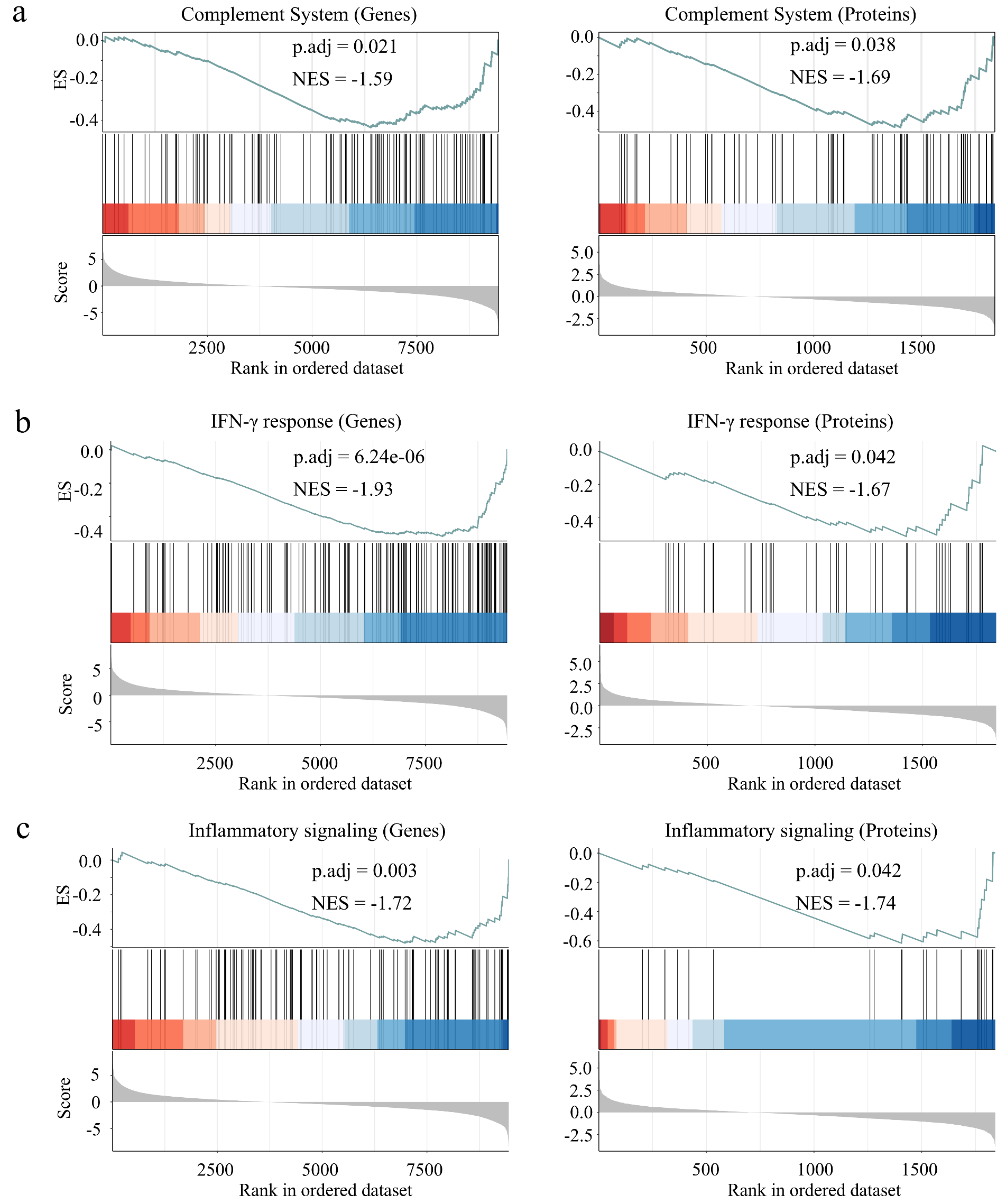

**Fig. 18. GSEA displays transcriptome and proteome pathways differentially enriched in treat vs. control groups including complement system (a), IFN-γ response (b), and inflammatory signaling (c).**

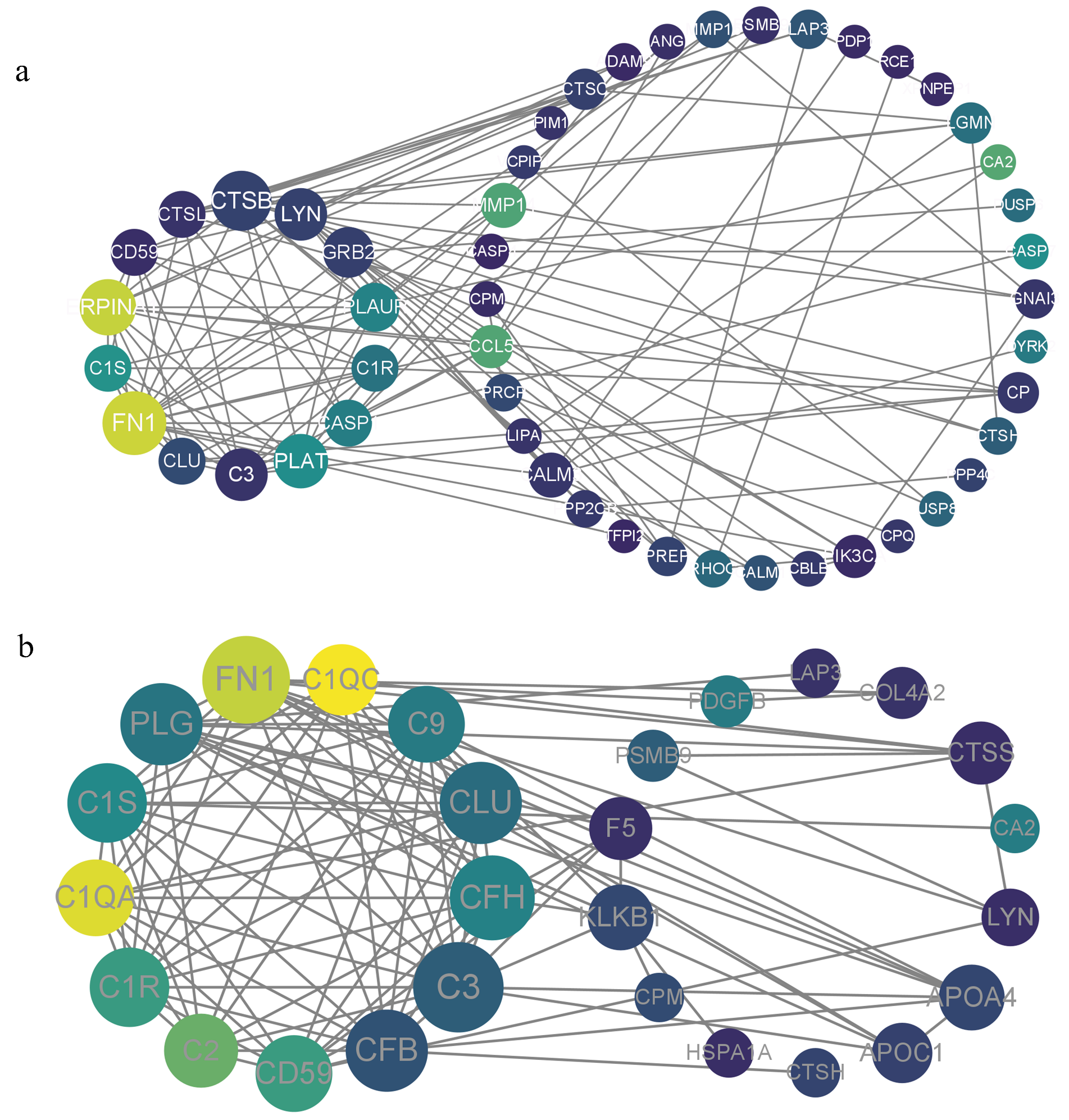

**Fig. 19. Protein-protein interaction networks of core complement components in the complement pathway by transcriptome (a) and proteome (b) analysis.**

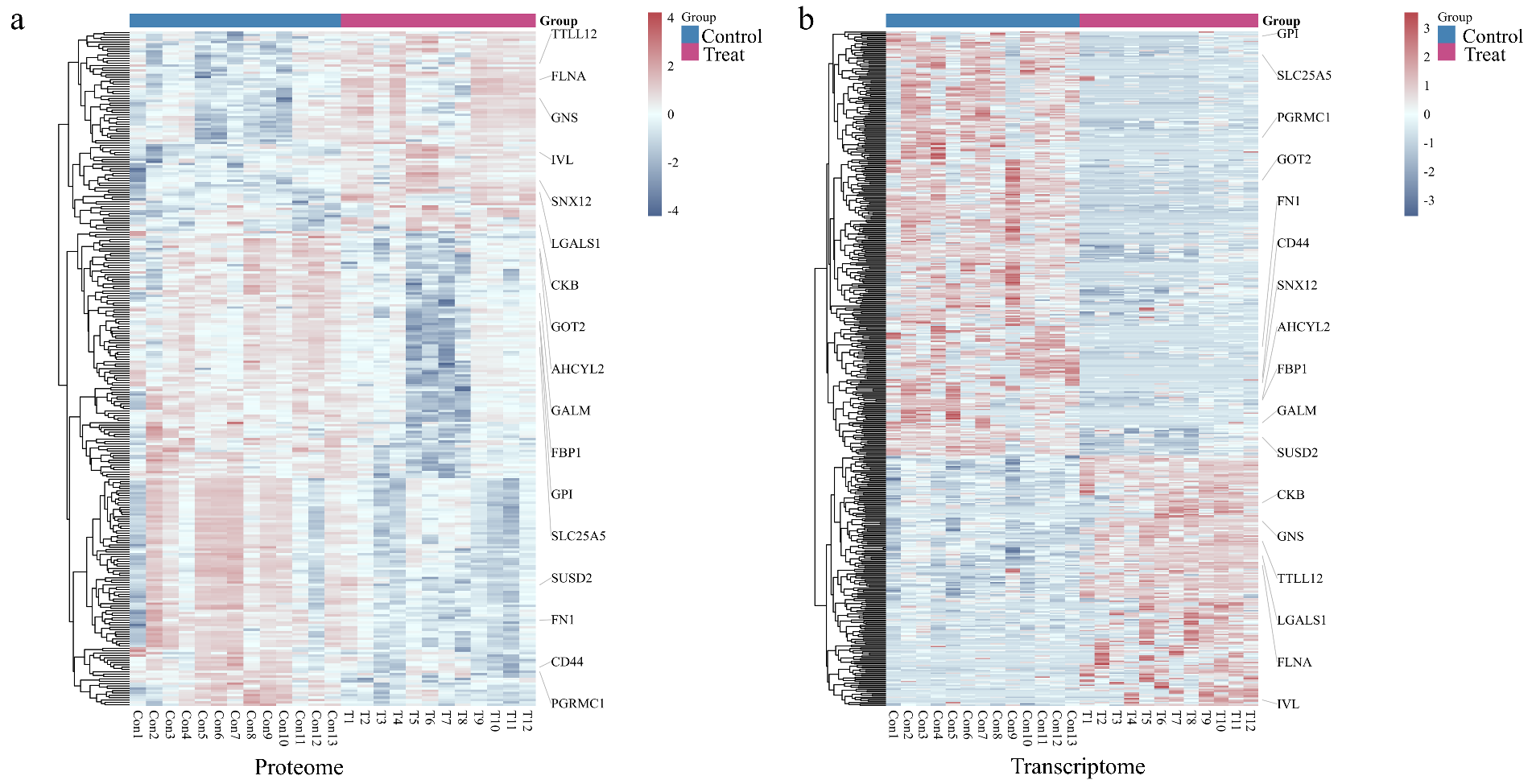

**Fig. 20. Heatmap of differential expression between proteome (a) and transcriptome (b).**

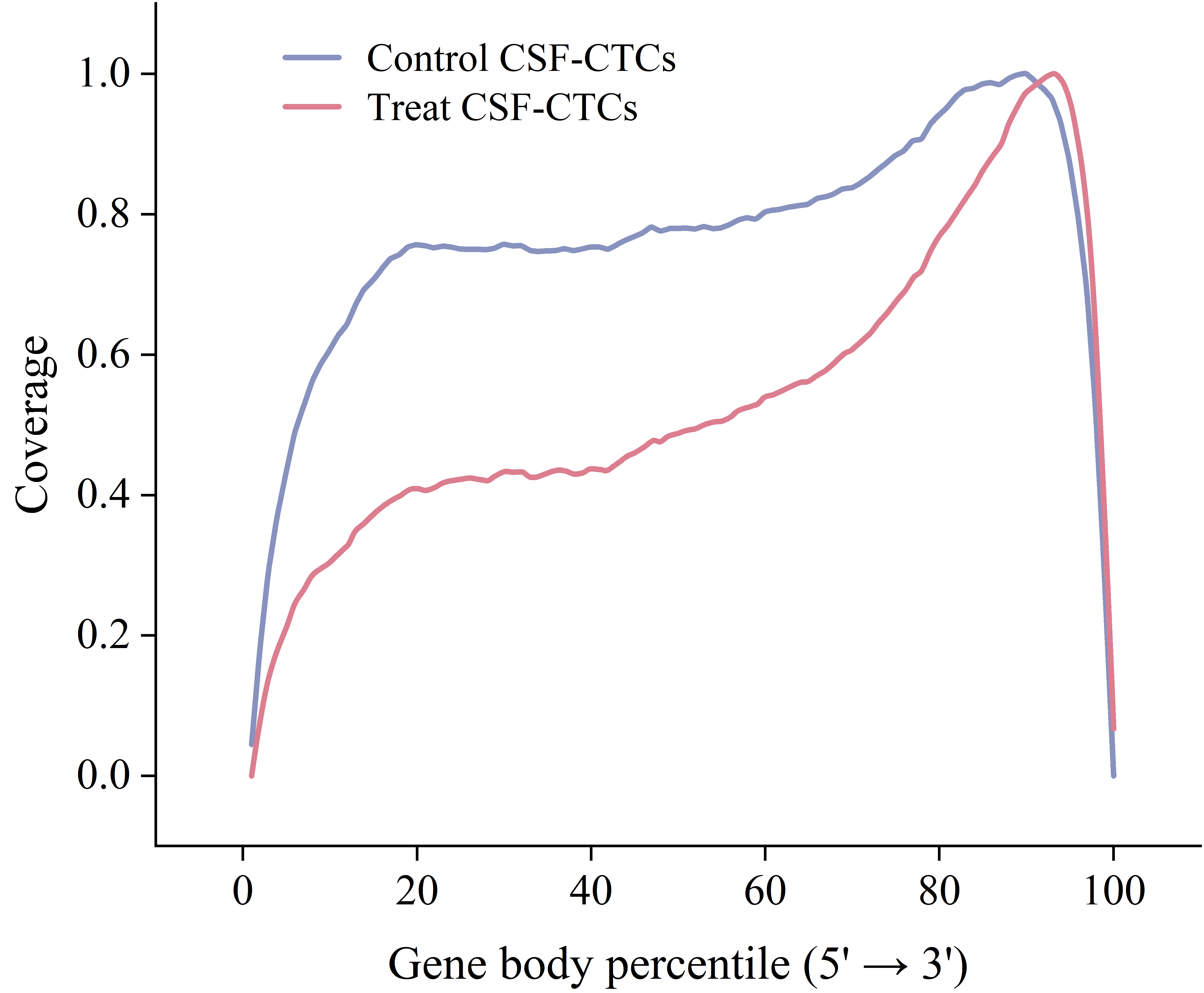

**Fig. 21. The sequencing gene body coverage ability of control CSF-CTCs and treat CSF-CTCs groups.**

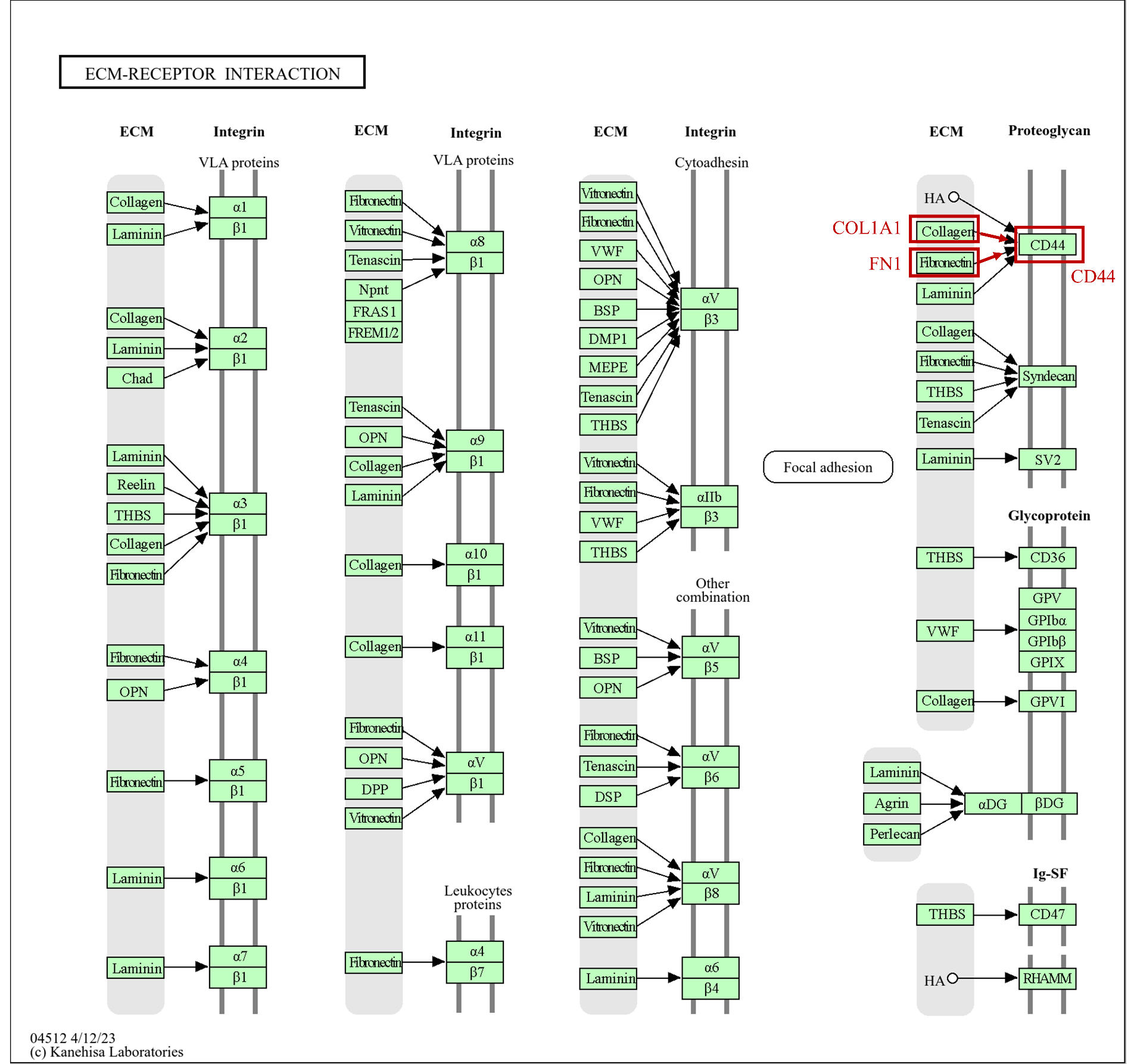

**Fig. 22. KEGG pathway of Ecm-Receptor Interaction including COL1A1, FN1, and CD44.**

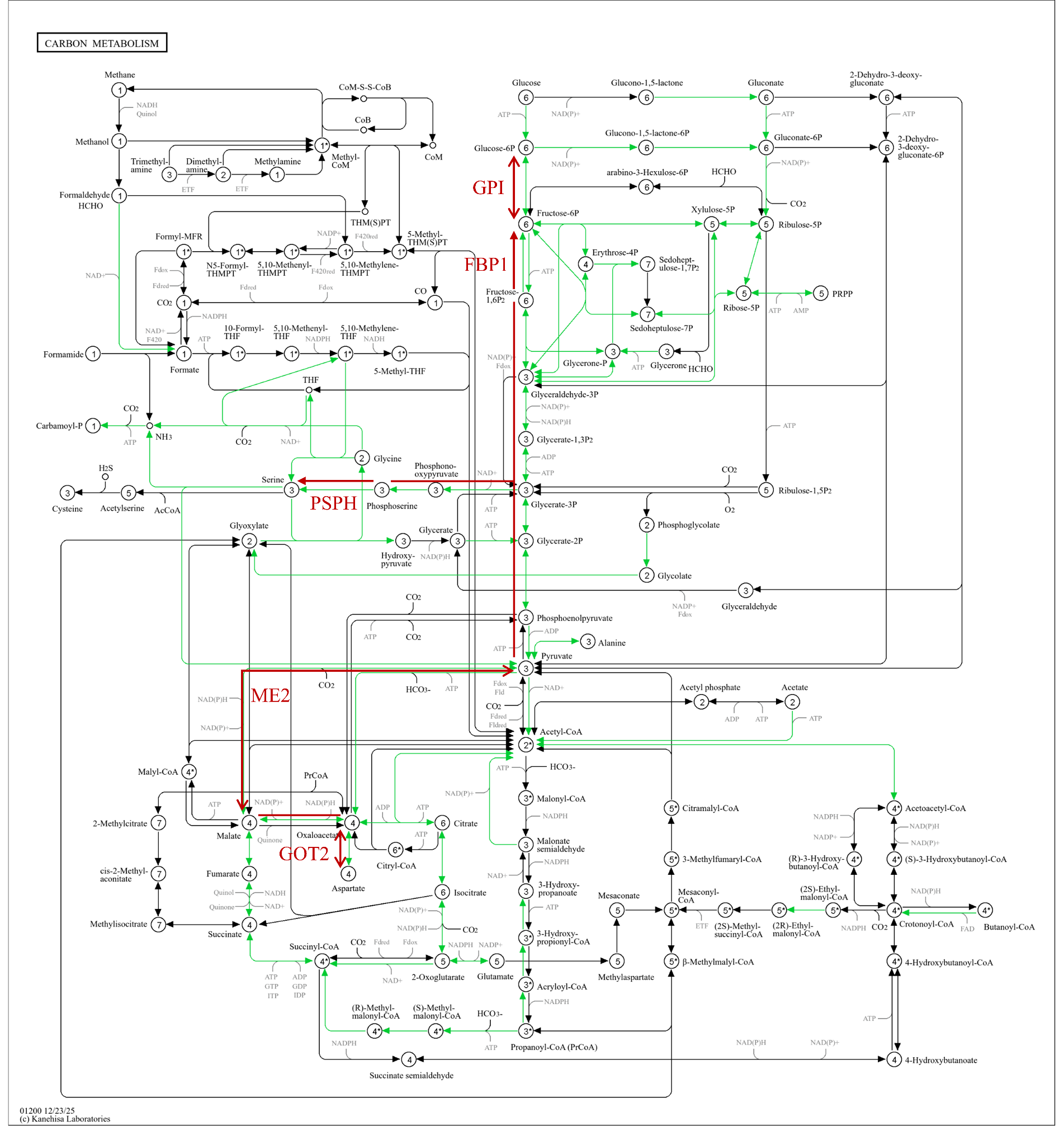

**Fig. 23. KEGG pathway of Carbon Metabolism including GPI, FBP1, PSPH, ME2, and GOT2.**

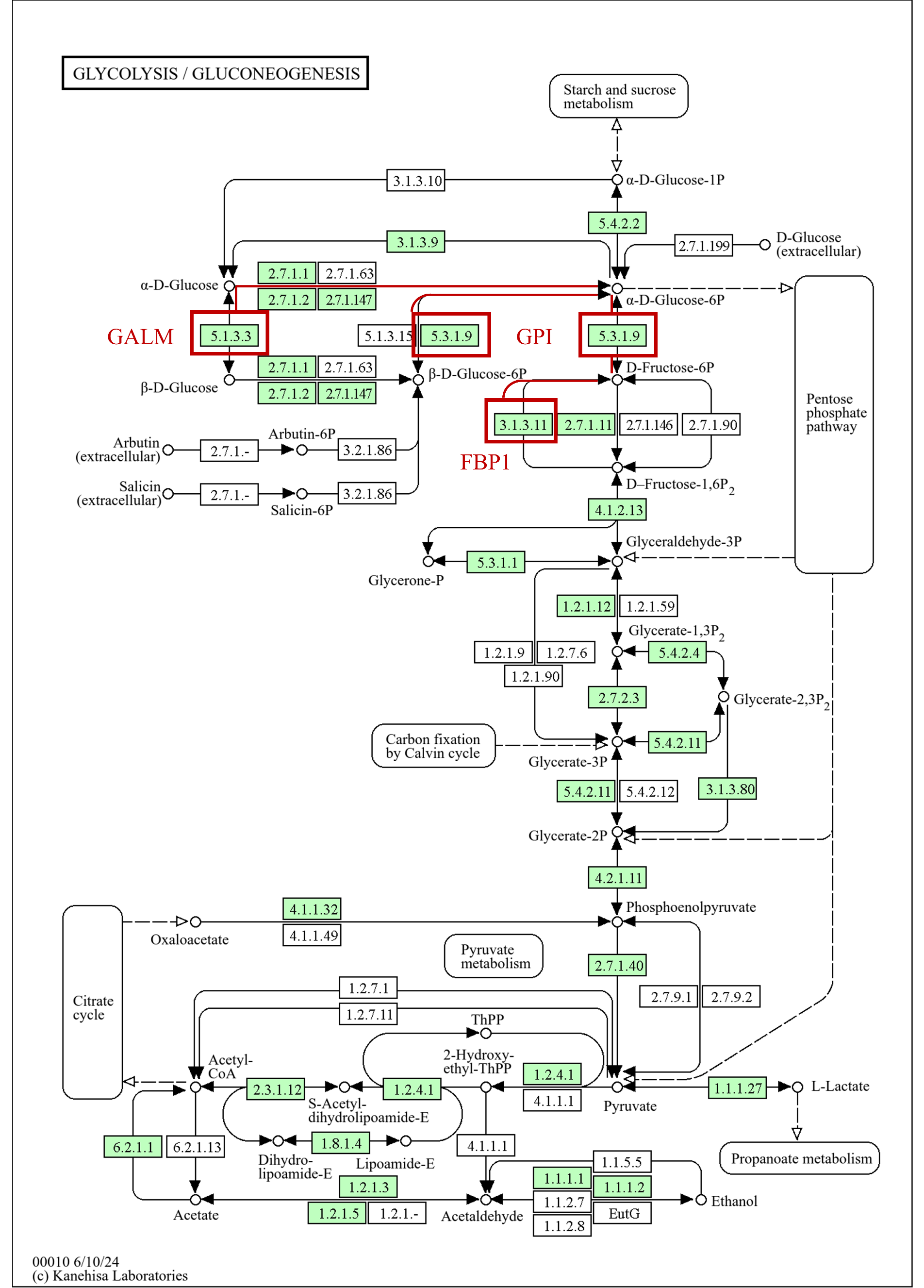

**Fig. 24. KEGG pathway of Glycolysis/Gluconeogenesis including GPI, FBP1, and GALM.**

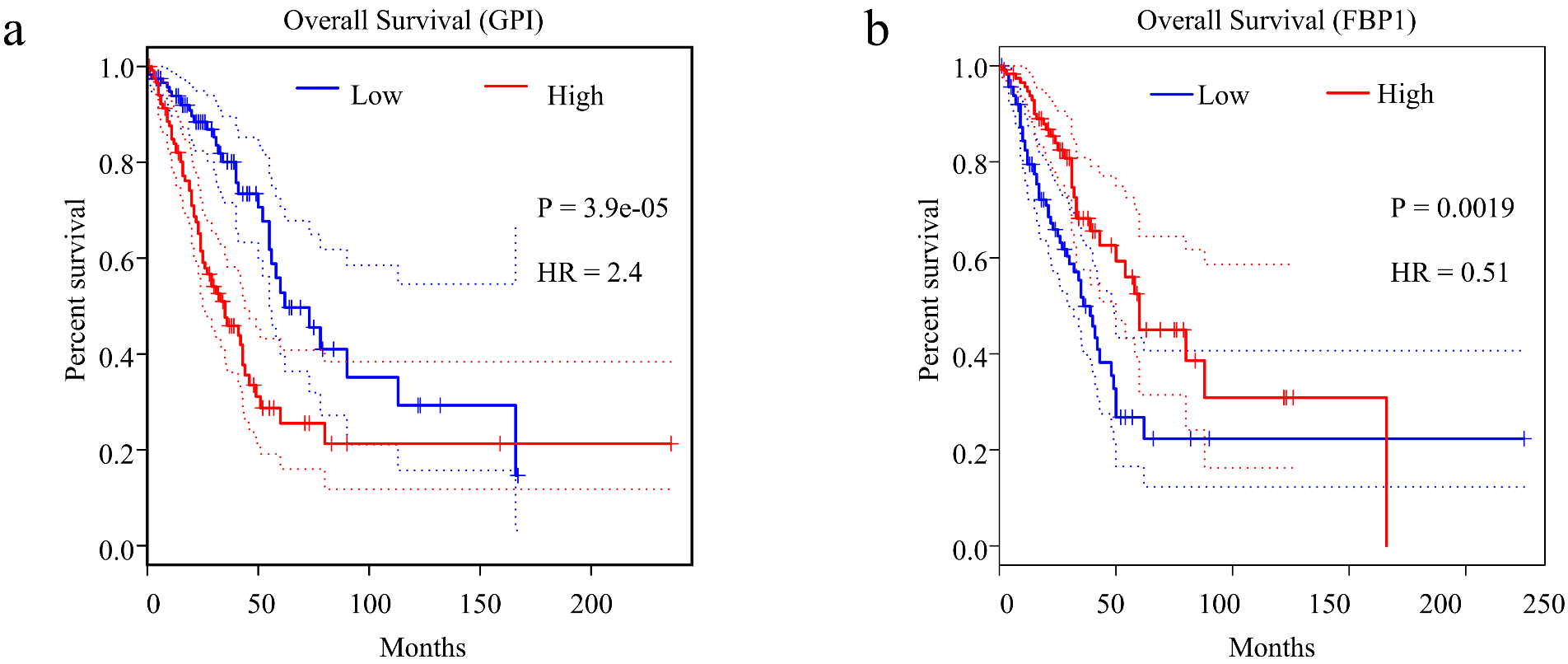

**Fig. 25. Kaplan-Meier overall survival analysis of the metabolic regulator GPI (a) and FBP1 (b) in the TCGA-LUAD cohort (n = 120).**

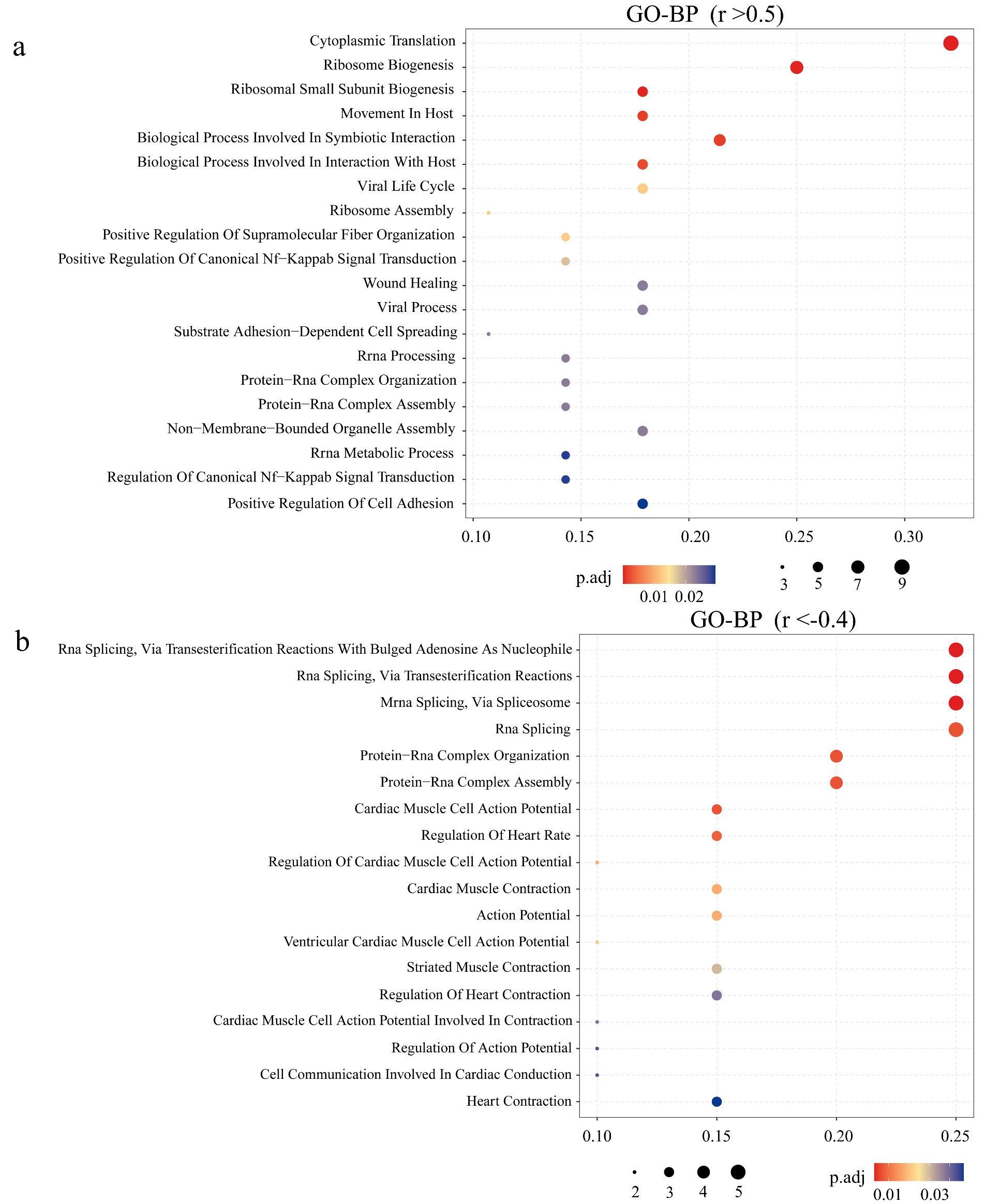

**Fig. 26. GO-Biological Process (GO-BP) analysis of positively correlated pairs (a, r > 0.5) and negatively correlated pairs (b, r < -0.4) in the multi-omics.**

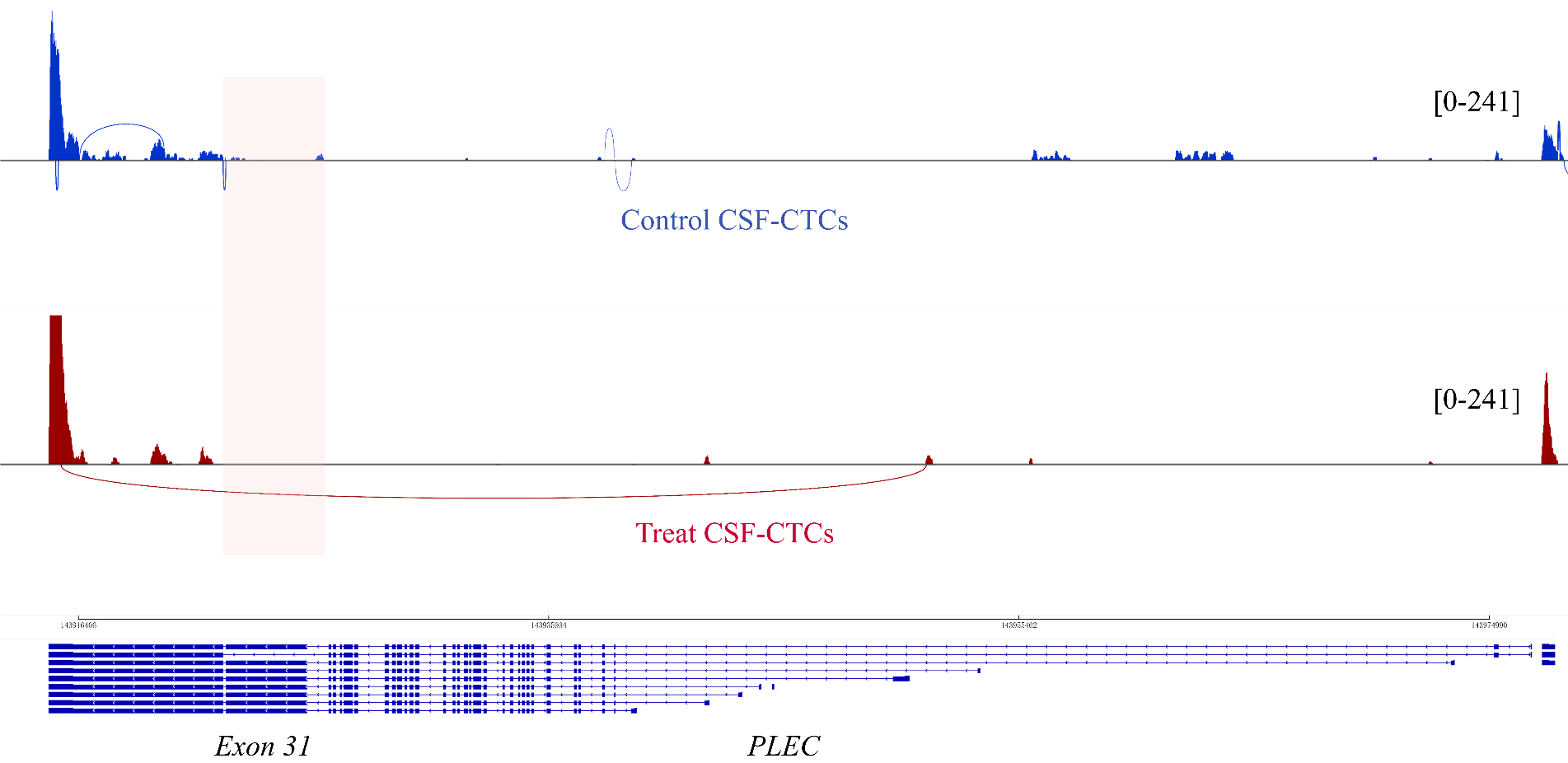

**Fig. 27. Sashimi plots visualization of RNA-seq reads mapping to the *PLEC* locus in control (top, blue) versus treat (bottom, red) CTCs.** Loops represent splice junctions, and the height of bars represents read coverage.

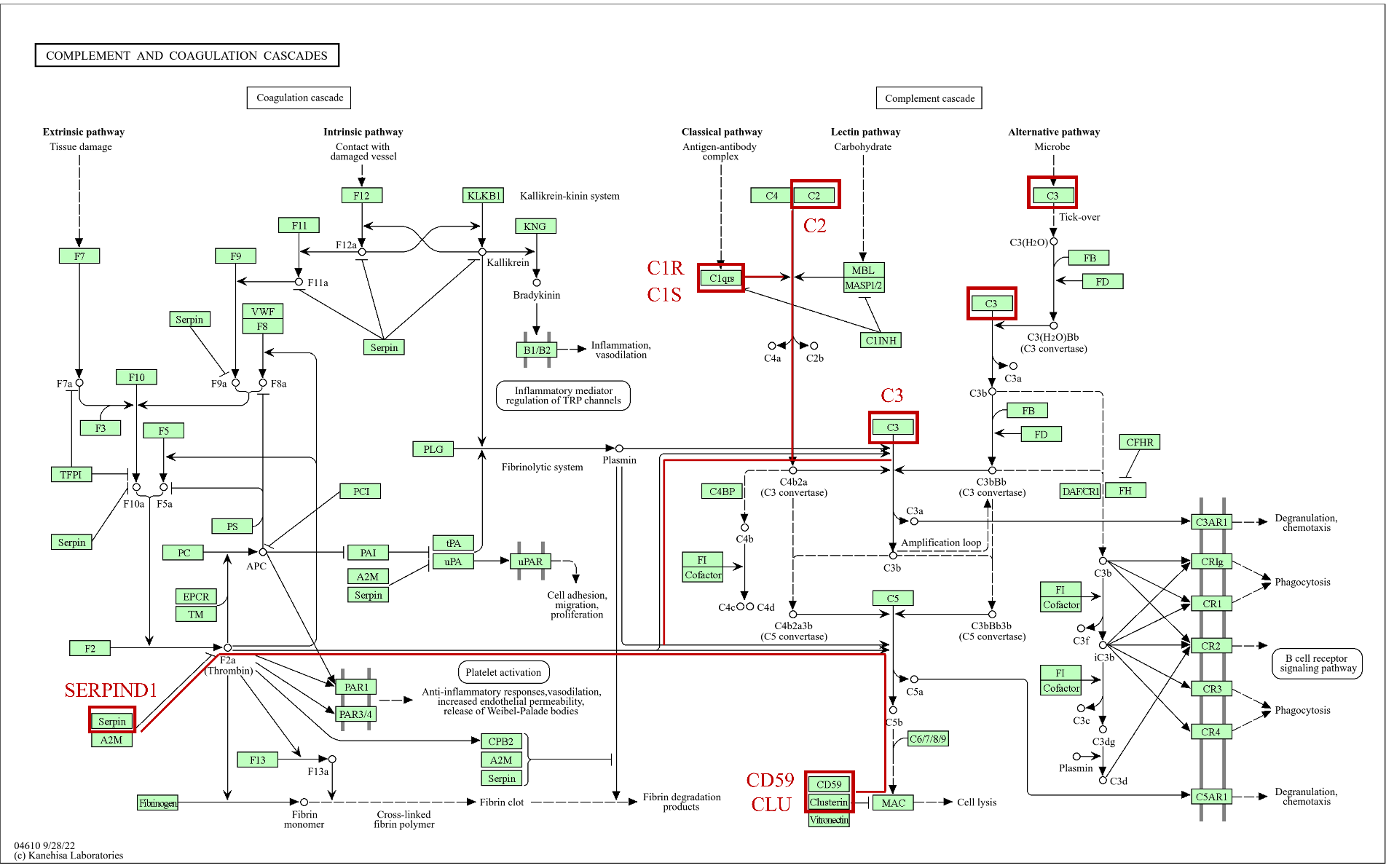

**Fig. 28. KEGG pathway of complement based on the Class III genes (concomitant downregulation at transcript and protein levels).**

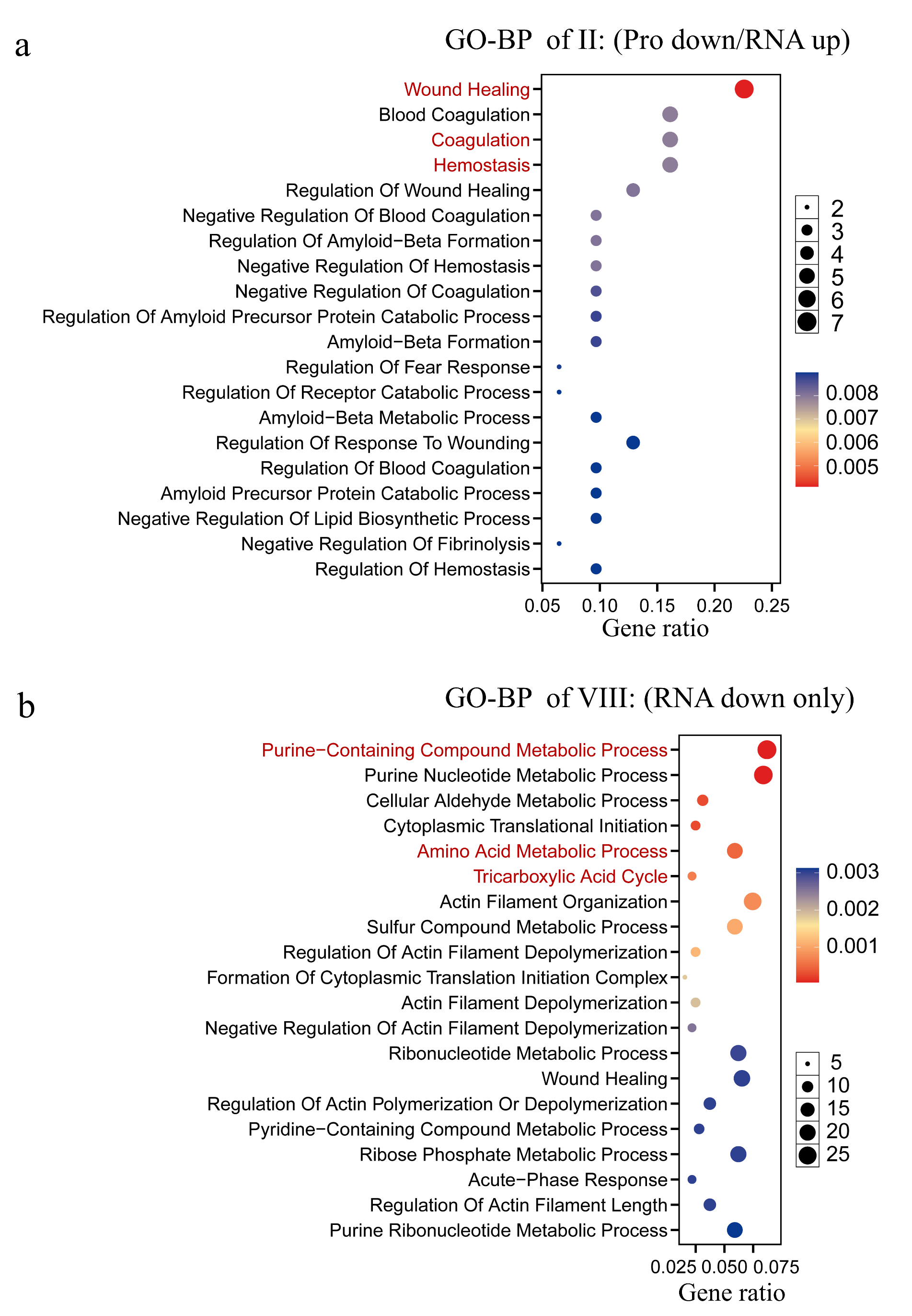

**Fig. 29. GO-BP analysis of protein down/RNA up (a) and RNA down only (b) in the multi-omics.**

**Table. 1. List of common differentially expressed gene-protein pairs**

| Gene ID | Protein log2FC | Protein adj.P | Gene log2FC | Gene adj.P |
| --- | --- | --- | --- | --- |
| AHCYL2 | -1.396266 | 0.008591 | -2.738499 | 0.015291 |
| AQP1 | -3.868766 | 0.000284 | -5.839413 | 0.000000 |
| BCAP31 | 1.574925 | 0.005698 | 1.731506 | 0.001445 |
| CA2 | -1.569493 | 0.021951 | -4.354793 | 0.037029 |
| CD44 | -2.112432 | 0.010519 | -5.975710 | 0.000011 |
| CDC42BPA | -0.874226 | 0.001918 | -2.295832 | 0.018513 |
| CKB | 1.289826 | 0.008970 | 2.403316 | 0.043817 |
| COL1A1 | -0.957512 | 0.035593 | 2.616185 | 0.032856 |
| COQ9 | -0.828869 | 0.048023 | -4.173946 | 0.018110 |
| CTSH | -1.078433 | 0.022367 | -2.226987 | 0.023146 |
| DNAJB6 | 0.611615 | 0.023057 | -2.562164 | 0.000305 |
| EMD | 2.163660 | 0.000932 | 2.510079 | 0.000066 |
| FAM177A1 | 2.855156 | 0.000160 | 1.264970 | 0.019813 |
| FBP1 | -2.931829 | 0.001291 | -3.328753 | 0.006776 |
| FLNA | 2.575336 | 0.000016 | 2.344520 | 0.000343 |
| FN1 | -2.679068 | 0.000539 | -6.240072 | 0.000001 |
| GALM | -1.263361 | 0.036479 | -4.992838 | 0.035891 |
| GNS | 1.600308 | 0.010230 | 2.329878 | 0.002854 |
| GOT2 | -1.434734 | 0.041232 | -2.485785 | 0.034035 |
| GPI | -2.117903 | 0.006704 | -2.509714 | 0.015693 |
| IVL | 6.380437 | 0.000000 | 6.962079 | 0.000001 |
| LGALS1 | 5.639534 | 0.000000 | 3.178643 | 0.001445 |
| MCCC2 | -1.001730 | 0.015959 | -4.540450 | 0.000607 |
| ME2 | -2.661375 | 0.000355 | -4.044503 | 0.017423 |
| NAPA | -0.933828 | 0.043779 | -4.213266 | 0.003817 |
| PGRMC1 | -1.691012 | 0.026941 | -3.483288 | 0.001033 |
| PIGR | -1.270966 | 0.037730 | -6.585336 | 0.000000 |
| PNPT1 | -0.753129 | 0.010040 | -3.911339 | 0.001498 |
| PPIA | 2.135797 | 0.000001 | 1.005818 | 0.019151 |
| PSPH | 1.097822 | 0.012062 | 3.729329 | 0.022557 |
| SLC25A5 | -2.027568 | 0.020400 | -2.155943 | 0.010655 |
| SNX12 | 2.007093 | 0.001450 | -6.021069 | 0.000087 |
| SUSD2 | -1.103254 | 0.004563 | -2.667009 | 0.033457 |
| TTLL12 | 1.519032 | 0.040056 | 3.626023 | 0.019174 |
| TXNL1 | -1.804848 | 0.002181 | -2.852395 | 0.039451 |
| UBA2 | -1.172209 | 0.038214 | -2.368594 | 0.019813 |
| ZNF185 | 1.086679 | 0.001024 | 4.529198 | 0.003757 |
